# PAIP1 couples mRNA export to cytoplasmic mRNP remodeling and poly(A) homeostasis

**DOI:** 10.64898/2026.07.31.742013

**Authors:** Daan Overwijn, Kerstin Dörner, Caroline de Almeida, Martin Müller, Aleksei Mironov, Robert Ivanek, Peter Skrinjar, Michelle Jennifer Gut, Isabel Solheim, Nicole Beuret, Thomas Bock, Matthias Zeug, Maciej Smialek, Sina Brandkamp, Timm Maier, Mihaela Zavolan, Claudia Isabelle Keller Valsecchi, Maria Hondele

## Abstract

Messenger ribonucleoproteins (mRNPs) acquire distinct protein compositions as they move from the nucleus to the cytoplasm, yet how this transition is coordinated and how inefficient remodeling affects downstream mRNA metabolism remains poorly understood. Here, we identify PAIP1 as a metazoan cofactor of the mRNA export ATPase DDX19. By binding a conserved N-terminal motif in DDX19, PAIP1 is recruited to the nuclear pore, where it promotes exchange of the nuclear poly(A)-binding protein PABPN1 for cytoplasmic PABPC1, thereby coupling mRNA export to cytoplasmic mRNP maturation. PAIP1 depletion alters cytoplasmic mRNP composition, mRNA stability and poly(A)-tail homeostasis. In Drosophila embryos, where poly(A)-tail regulation of maternal mRNAs directs early development, maternal PAIP1 depletion shortens poly(A)-tails, delays zygotic genome activation, and causes severe developmental defects. Our findings identify an export-coupled mRNP maturation pathway linking PABP exchange to downstream mRNA metabolism.

## INTRODUCTION

In eukaryotes, mRNA production in the nucleus is spatially separated from translation in the cytoplasm. Newly synthesized mRNAs leave the nucleus as messenger ribonucleoprotein particles (mRNPs) bound by a characteristic set of nuclear RNA-binding proteins. Upon entry into the cytoplasm, mRNPs undergo extensive remodeling as nuclear factors are replaced by cytoplasmic proteins that regulate translation, storage, and decay^1,2^. A central event in this transition is the exchange of nuclear poly(A)-binding proteins (PABPs), predominantly PABPN1, for cytoplasmic PABPs, predominantly PABPC1. PABPN1 supports nuclear poly(A)-tail elongation and quality control^3–5^, whereas PABPC1 promotes translation and influences mRNA stability in the cytoplasm^6–8^. Replacement of PABPN1 by PABPC1 therefore constitutes a key remodeling step in the establishment of the cytoplasmic mRNP state. However, how this transition is coordinated and whether its efficiency has functional consequences for newly exported mRNPs remain poorly understood.

Newly exported mRNPs enter the cytoplasm through the nuclear pore complex (NPC), making the cytoplasmic face of the pore a plausible site for coordinating the transition from nuclear to cytoplasmic mRNPs. Here, the DEAD-box ATPase DDX19 (DDX19A and DDX19B in humans, collectively referred to here as DDX19; Dbp5 in yeast) drives directional mRNA export^9–13^. Several mechanisms have been proposed to mediate PABP exchange, including export-coupled remodeling by DDX19/Dbp5, intrinsic differences in the RNA-binding properties of PABPN1 and PABPC, and the pioneer round of translation^11,13–16^. The relative contribution of these pathways to PABP exchange remains unresolved, in part because they also perform essential functions in mRNA export or downstream cytoplasmic gene expression, making PABP exchange itself difficult to perturb selectively.

Here, we identify polyadenylate-binding protein-interacting protein 1 (PAIP1) as a DDX19 interactor required for the establishment of a mature cytoplasmic mRNP state. PAIP1 positions PABPC1 at NPC-proximal sites and promotes PABPN1-to-PABPC1 exchange. Loss of PAIP1 alters mRNP composition and disrupts transcript abundance, poly(A)-tail homeostasis and mRNA stability in a transcript-specific manner, preferentially altering poly(A)-tail length of poorly translated mRNAs while destabilizing long-lived transcripts. During early *Drosophila* development, where gene expression depends predominantly on post-transcriptional regulation of maternally deposited mRNAs, maternal PAIP1 depletion causes widespread poly(A)-tail shortening, delayed zygotic genome activation and severe embryonic development defects. Together, these findings identify an export-coupled mRNP maturation pathway that links PABP exchange to downstream mRNA metabolism and developmental gene regulation.

## RESULTS

### PAIP1 is a shared interactor of PABPC1 and DDX19

To identify factors that might promote the PABPN1-to-PABPC1 transition, we searched for proteins that connect PABPC1 to the mRNA export machinery, particularly DDX19. We combined two complementary approaches. First, we intersected the BioGRID interactomes of PABPC1 and DDX19A/B^17^, identifying 30 shared interactors (criterion 1, (Figure 1A)). In parallel, we used AlphaFold-Multimer^18^ to predict direct interactions between DDX19 and its reported BioGRID interactors (n = 251 for DDX19A/B), which recovered several established DDX19 cofactors, including NUP214, GLE1, CTIF and MIF4GD/SLIP1, as positive controls (criterion 2, (Figures 1B and S1A))^12,19–21^. Only two proteins satisfied both criteria: GLE1 and Poly(A)-binding protein– Interacting Protein 1 (PAIP1). Because GLE1 is not predicted to be a direct interactor of PABPC1 (Figure S1B), we focused on PAIP1, a poorly characterized cytoplasmic PABPC cofactor with no previously reported role in mRNA export.

**Figure 1.**
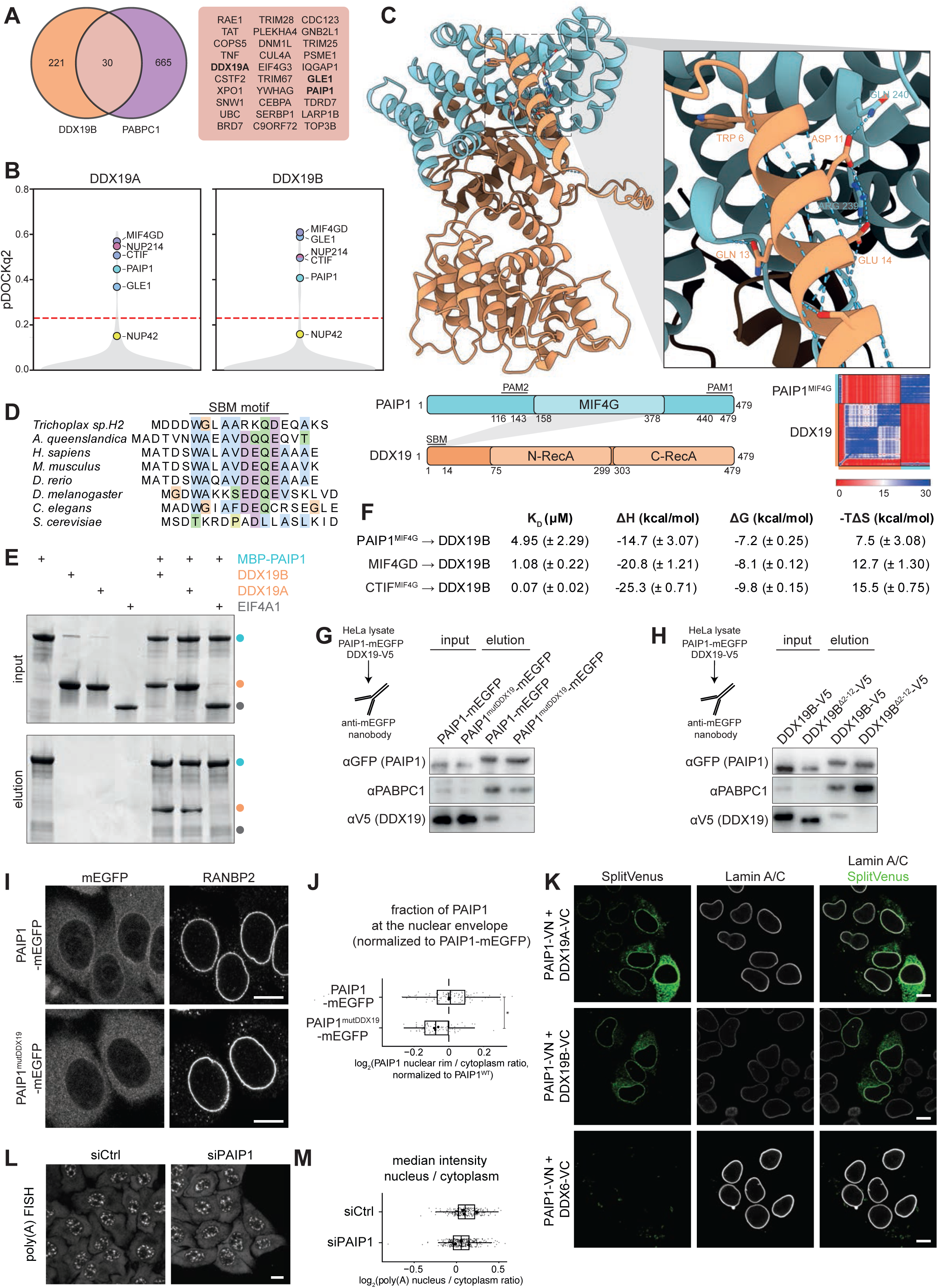
PAIP1 is a conserved DDX19 cofactor that binds the N-terminal SBM through its MIF4G domain. (A) Overlap between BioGRID-curated DDX19 and PABPC1 interactomes. Shared cofactors are listed. (B) AlphaFold-Multimer screen of DDX19A/B against BioGRID interactors. Predicted complexes are ranked by pDockQ2 score. The reliability threshold (pDockQ2 ≥ 0.23) is indicated.Known cofactors are highlighted. (C) AlphaFold-Multimer model of the PAIP1 MIF4G domain bound to DDX19; domain architecture of PAIP1 and DDX19; predicted aligned error (PAE) plot of alphafold multimer model between PAIP1 and the conserved N-terminal SBM motif of DDX19. (D) Conservation of the DDX19 SBM motif across orthologs. (E) In vitro pull-down of recombinant DDX19A/B or EIF4A1 with bead-immobilized MBP–PAIP1. (F) Summary of isothermal titration calorimetry (ITC) measurements for PAIP1, MIF4GD/SLIP1 or CTIF titrated into DDX19B. Values represent mean ± SD, N = 2. (G, H) Stable HeLa cell lines expressing siRNA-resistant PAIP1-mEGFP or PAIP1^mutDDX19^-mEGFP at endogenous level were transiently transfected to express DDX19B-V5 or DDX19^Δ2–12^-V5 for 24h. GFP immunoprecipitates were analyzed by immunoblotting with the indicated antibodies. N = 3. (I) Stable HeLa cell lines expressing PAIP1-mEGFP or PAIP1^mutDDX19^-mEGFP near endogenous level were fixed and immunostained with an antibody against RANBP2. Scale bar: 10 µm, N = 3, n ≥ 58 cells. (J) Enrichment of PAIP1 at the nuclear rim, calculated for each cell as the ratio of median PAIP1 fluorescence intensity at the nuclear rim to median cytoplasmic intensity. Ratios were normalized within each biological replicate to the mean ratio of PAIP1^WT^–mEGFP cells and are displayed on a log₂ scale. Black dots indicate the log₂-transformed mean normalized ratio for each biological replicate. Two-tailed one-sample t-test on the log₂-transformed ratios from three biological replicates against 0; p = 0.0482 (*). (K) Split-Venus bimolecular fluorescence complementation (BiFC) analysis in HeLa cells transiently co-transfected with Venus N-terminus– tagged PAIP1 (PAIP1-VN) and Venus C-terminus–tagged DDX19A/B (DDX19A/B-VC) or DDX6-V5-VC; cells were co-stained for LaminA/C. Scale bar: 10 µm, N = 3, n ≥ 32 cells. (L) HeLa cells treated with control siRNA (siCtrl) or siRNA targeting PAIP1 were analyzed by poly(A) RNA FISH. N = 3, n ≥ 100 cells. (M) Poly(A) RNA distribution was quantified for individual cells as the ratio of median nuclear to median cytoplasmic poly(A) intensity and is displayed on a log₂ scale. Small grey dots represent individual cells and large black dots indicate the log₂-transformed mean nucleus/cytoplasm ratio for each biological replicate. Two-tailed one-sample t-test on the log₂ - transformed siPAIP1/siCNTL mean ratios from three biological replicates against 0; *p* = 0.385 (ns).

### PAIP1 engages the conserved DDX19 SBM through its MIF4G domain

PAIP1 is a translation co-activator^22–24^ that binds PABPC through two conserved PABP-interacting motifs, PAM1 and PAM2, like its paralogue PAIP2^25,26^. Unlike PAIP2, PAIP1 additionally contains a central MIF4G domain, a structural fold found in several cofactors of DEAD-box ATPases^24,27^. Inspection of the AlphaFold-Multimer models (Figures 1C and S1C-E) revealed that the PAIP1 MIF4G domain engages an evolutionarily conserved ∼12-residue segment at the extreme N-terminus of DDX19. This region encompasses the previously described SLIP1-Binding Motif (SBM, DDX19 residues 6-14), which serves as docking site for the MIF4G-domain proteins CTIF and MIF4GD/SLIP1, factors involved in specialized translation pathways (histone and pioneer translation, respectively)^20,21^ (Figure 1D).

In the PAIP1-DDX19 model, the SBM motif adopts an α-helix that binds a groove in the PAIP1 MIF4G domain, closely resembling the SBM-CTIF or SBM-MIF4GD complexes^20,21^. This interaction mode is structurally and functionally distinct from the canonical interaction of MIF4G domains with the DEAD-box ATPase RecA core, which is utilized e.g. by the GLE1 MIF4G domain to stimulate DDX19 ATPase activity during mRNA export^28–30^. Notably, the PAIP1 SBM-binding groove is also targeted by the SARS-CoV-2 protein NSP3 (Figure S1F)^31^. Our findings identify PAIP1 as a previously unrecognized SBM-binding protein, linking DDX19 not only to specialized translation pathways but also to the canonical PABPC / poly(A)-dependent translation machinery.

To determine when the DDX19-PAIP1 interaction emerged during evolution, we compared the conservation of the core mRNA export factors DDX19 and GLE1, the SBM motif, and SBM-binding MIF4G proteins across eukaryotes. While DDX19 and GLE1 are deeply conserved and present in fungi, the SBM motif and its MIF4G-binding partners (CTIF, MIF4GD/SLIP1 and PAIP1) are restricted to metazoans (Figures 1D and S1G-H). PAIP1 and the predicted PAIP1-DDX19 interface are already present in early metazoans, including placozoans such as *Trichoplax*, and sponges such as *Amphimedon queenslandica* (Figure S1I), suggesting that SBM-mediated PAIP1 recruitment emerged early during metazoan evolution.

### PAIP1 binds DDX19 without modulating ATPase activity

We next validated the predicted PAIP1–DDX19 interaction experimentally. In vitro pull-down assays with recombinant proteins verified that maltose binding protein (MBP)-tagged full-length PAIP1 robustly bound DDX19A and DDX19B, but not EIF4A1, a DEAD-box ATPase previously proposed to engage the PAIP1 MIF4G domain^22,32^ (Figure 1E). Isothermal titration calorimetry (ITC) further showed that the PAIP1 MIF4G domain binds full-length DDX19B with low-micromolar affinity (K_D_ ≈ 4.95 µM). This affinity is comparable to that of MIF4GD/SLIP1 (K_D_ ≈ 1.08 µM) but substantially weaker than the affinity of the CTIF MIF4G domain (K_D_ ≈ 69 nM) (Figures 1F and S1J). Quantitative proteomics datasets from cell lines and tissues^33–36^ however indicate that PAIP1 is approximately 4–37-fold more abundant than CTIF or MIF4GD/SLIP1, and total DDX19A/B generally exceeds the abundance of all three SBM-binding proteins combined (Supplementary table 1). Thus, despite its lower intrinsic affinity, PAIP1 should efficiently engage the DDX19 SBM in cells.

To test whether the predicted SBM–MIF4G interface mediates binding in cells, we generated complementary mutations in both proteins. Deletion of DDX19B residues 2–12 (DDX19B^Δ2–12^) removed the SBM, whereas substitution of two residues within the predicted SBM-binding groove of PAIP1 (R239A/Q240A, PAIP1^mutDDX19^) targeted the reciprocal interface (Figure 1C). In stable HeLa cells expressing PAIP1–mEGFP variants, wild-type PAIP1 efficiently recovered transiently transfected DDX19B–V5, whereas PAIP1^mutDDX19^ showed markedly reduced DDX19 co-precipitation while retaining PABPC1 binding (Figure 1G). Conversely, wild-type PAIP1 failed to recover DDX19B^Δ2–12–V5^ (Figure 1H). Proteomic analysis of V5 immunoprecipitates from stable DDX19B–V5 cell lines likewise showed that endogenous PAIP1 was enriched with wild-type DDX19B but not with DDX19B^Δ2–12^ (Figure S1K).

Because MIF4G-domain proteins frequently regulate the enzymatic activity of DEAD-box ATPases, we tested whether PAIP1 alters DDX19 ATPase activity in vitro. However, recombinant PAIP1 had no detectable effect on RNA-stimulated DDX19 ATPase activity (Figure S2A), consistent with its engagement of the N-terminal SBM motif rather than the helicase core.

Together, these findings establish PAIP1 as a bona fide DDX19 interactor that binds the N-terminal SBM without regulating ATPase activity, suggesting that the primary function of the interaction is to recruit PAIP1 to DDX19-containing mRNP export complexes.

### DDX19 recruits PAIP1 to NPC-proximal sites

If DDX19 recruits PAIP1 to the mRNA export machinery, at least a fraction of PAIP1 should localize to the cytoplasmic face of the NPC. Because available antibodies did not reliably detect endogenous PAIP1 by immunofluorescence, we analyzed stable HeLa cell lines expressing PAIP1–mEGFP at near-endogenous levels (Fig S2B). PAIP1–mEGFP was predominantly cytoplasmic but showed modest enrichment at the nuclear rim, where it colocalized with the NPC marker RANBP2 (Figure 1I). This enrichment was lost for PAIP1^mutDDX19^–mEGFP (Figures 1I and 1J), indicating that SBM-dependent docking to DDX19 recruits PAIP1 to NPC-proximal sites.

To determine whether nuclear-rim associated PAIP1 interacts with DDX19, we used Split-Venus bimolecular fluorescence complementation (BiFC). In this assay, candidate proteins are fused to complementary N- and C-terminal fragments of the Venus fluorescent protein (VN and VC), with fluorescence reconstituted only when the two partners come into close proximity (Figure S2C)^37^. Co-expression of PAIP1–VN with DDX19A/B–VC produced a prominent signal at the nuclear rim (marked by LaminA/C immunofluorescence), together with weaker diffuse cytoplasmic fluorescence (Figure 1K). Pairing DDX19-VC with GLE1-VN, CTIF–VN, or MIF4GD/SLIP1–VN yielded a similarly rim-enriched pattern (Figures S2D and S2E), consistent with recruitment of all four MIF4G cofactors to DDX19 at or near NPCs. By contrast, PAIP1–VN produced no signal with the unrelated cytoplasmic helicase DDX6–VC (Figure 1K). Thus, PAIP1 associates with DDX19 at or near the NPC.

### PAIP1 promotes PABPC1 association with DDX19 and the NPC

Although PAIP1 did not regulate DDX19 ATPase activity, its recruitment to DDX19 and the NPC suggested a role in an mRNA-export-coupled process. We first asked whether the PAIP1-DDX19 interaction is required for bulk poly(A)+ mRNA export. As expected, depletion of DDX19A/B caused strong nuclear retention of poly(A)+ RNA. This phenotype was rescued equally well by wild-type DDX19A/B and the PAIP1-binding deficient DDX19A/B^Δ2–12^ mutant (Figures S2F-H), indicating that SBM-dependent PAIP1 recruitment is dispensable for bulk export. PAIP1 depletion likewise did not cause nuclear accumulation of poly(A)+ RNA (Figures 1L, 1M and S2I). The PAIP1–DDX19 interaction might therefore function primarily downstream of mRNA export through the NPC.

Given that PAIP1 also binds the cytoplasmic poly(A)-binding protein PABPC1, we hypothesized that PAIP1 could promote PABPC1 loading onto newly exported mRNPs. Such a mechanism would require simultaneous engagement of DDX19 and PABPC1. Indeed, AlphaFold-Multimer models support a ternary arrangement in which the PAIP1 MIF4G domain binds DDX19, while the N-terminal PAM2 motif and C-terminal PAM1-containing region engage PABPC1 (Figures 2A, 2B and S3A-D). These predicted contacts agree with previous biochemical and structural studies of PAIP1-PABPC1 complexes^22,26,38^. Consistent with this model, recombinant MBP-PABPC1 efficiently pulled down DDX19 only in the presence of PAIP1, whereas no interaction between PABPC1 and DDX19 was detected in the absence of PAIP1 (Figures S3E), demonstrating that PAIP1 is sufficient to bridge both proteins in vitro. Size-exclusion chromatography coupled to multi-angle light scattering (SEC-MALS) further showed that PAIP1 forms homodimers (Figures S3F and S3G), raising the possibility that one PAIP1 dimer recruits two PABPC molecules to DDX19.

**Figure 2.**
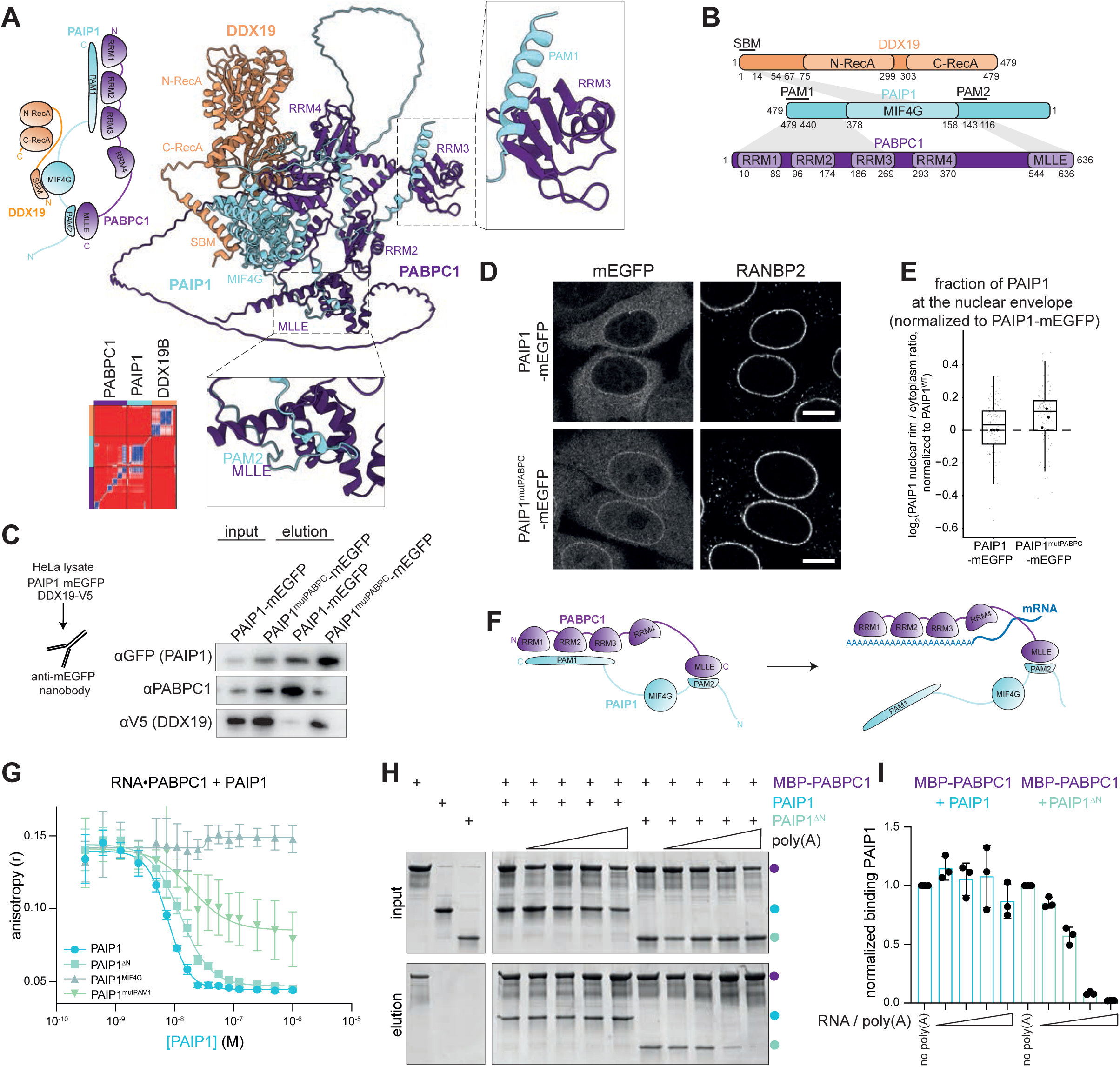
PAIP1 recruits PABPC1 to DDX19 at the NPC. (A) AlphaFold-Multimer model of full-length PAIP1, DDX19 and PABPC1 and corresponding PAE plots. (B) Schematic representation of interaction interfaces between DDX19, PAIP1 and PABPC1. (C) Stable HeLa cell lines expressing PAIP1–mEGFP or PAIP1^mutPABPC^-mEGFP at endogenous level were transiently transfected with DDX19-V5 for 48 h, followed by GFP immunoprecipitation and immunoblotting with the indicated antibodies. N = 3. (D) HeLa cell lines stably expressing PAIP1-mEGFP or PAIP1^mutPABPC^-mEGFP at endogenous level were fixed and immunostained with RANBP2. Scale bars 10 µm, N = 3, n ≥ 58 cells. (E) Fraction of PAIP1 at the nuclear rim in Figure 2D, calculated for each cell as the ratio of median PAIP1 fluorescence intensity at the nuclear rim to median cytoplasmic intensity. Ratios were normalized within each biological replicate to the mean ratio of PAIP1^WT^–mEGFP cells and are displayed on a log₂ scale. Small grey dots represent individual cells; large black dots indicate the log₂-transformed mean normalized ratio for each biological replicate. Two-tailed one-sample t-test on the log₂-transformed ratios from three biological replicates against 0; p=0.157 (ns). (F) Schematic of the interactions between PAIP1, PABPC1, and poly(A) RNA, illustrating competition between PAIP1 PAM1 and poly(A) for binding to PABPC1 RRMs 1-3. (G) Fluorescence polarization competition assay in which recombinant PAIP1 variants were titrated against a fixed concentration of a preformed A18 RNA–PABPC1 complex (see Figure S3G), N = 3. (H) Pull-down assays testing poly(A) RNA-dependent dissociation of PAIP1 or PAIP1^ΔN^ from immobilized MBP-PABPC1. N = 2. (I) Quantification of (J). Mean ± SD, N = 3.

To disrupt PABPC binding, we combined point substitutions in the PAM1-containing region (Y466A/F469A/D459A/Q420A) with deletion of PAM2, yielding PAIP1^mutPABPC^ (Figure S3A), and generated a stable PAIP1^mutPABPC^–mEGFP cell line. Immunoprecipitation followed by Western blotting or mass spectrometry confirmed that while PAIP1^WT^-mEGFP efficiently recovered endogenous PABPC proteins, these interactions were strongly reduced for PAIP1^mutPABPC^-mEGFP (Figures 2C and S3H). Strikingly, PAIP1^mutPABPC^–mEGFP accumulated in the nucleus and at the nuclear rim relative to PAIP1^WT^-mEGFP (Figures 2D, 2E and S2B), accompanied by increased recovery of nuclear proteins in IP–MS (Figure S3H). These observations suggest that PAIP1 might undergo nucleocytoplasmic shuttling, and that interaction with PABPC promotes its release from the NPC-proximal sites, potentially within a complex that accompanies newly exported mRNPs into the cytoplasm.

### PAIP1 uses distinct interaction interfaces to remain associated with PABPC1 during poly(A)-tail binding

For PAIP1 to deliver PABPC1 to newly exported mRNPs, their interaction must remain compatible with poly(A)-tail binding. AlphaFold-Multimer predicts that PAIP1 contacts PABPC1 at two sites: PAIP1 PAM1 engages the RNA-binding RRMs (RRM1–3) of PABPC1, whereas PAM2 binds the distal MLLE/PABC domain (Figure 2F). This architecture suggested that PABPC1 could simultaneously engage the poly(A) tail and PAIP1; even if poly(A) RNA displaces PAM1 from the RRMs, the PAM2–MLLE interaction would maintain the PAIP1–PABPC1 complex.

To determine whether PAM1 competes with RNA for PABPC1 RRM binding, we performed fluorescence-polarization assays. Recombinant PABPC1 bound fluorescently labelled A_18_ RNA with an apparent K_D_ of ∼ 0.1 nM, consistent with the low nanomolar affinity previously reported^14,39^ (Figure S3I). Addition of full-length PAIP1 displaced PABPC1 from pre-formed PABPC1–A_18_ RNA complexes with an apparent IC₅₀ of ∼ 8 nM (Figure 2G), indicating that the predicted interaction of PAIP1 PAM1 with PABPC1 RRMs does occur in solution and competes with RNA for the similar surfaces. As expected, a construct lacking the MLLE-binding PAM2 motif (PAIP1^ΔN^) displaced PABPC1 from RNA with similar efficiency (IC₅₀ ∼ 13 nM). By contrast, constructs with mutated PAM1 (PAIP1^mutPAPM1^) or the isolated MIF4G domain (which lacks both PAM1 and PAM2) significantly reduced (IC₅₀ ∼ 19.4 nM) resp. completely abolished PABPC1 displacement from RNA. Thus, the PAM1-containing region and poly(A) RNA engage overlapping surfaces on the PABPC1 RRM domains.

We then tested whether PAIP1 can remain bound to PABPC1 via their second PAM2–MLLE interface even when RNA blocks the PABPC1 RRMs. For this, we immobilized recombinant MBP–PABPC1 on beads and examined PAIP1 binding in the presence of increasing concentrations of synthetic poly(A) RNA. Full-length PAIP1 remained stably associated with PABPC1 even at high poly(A) concentrations (Figures 2H and 2I). By contrast, PAIP1^ΔN^, which lacks PAM2 and therefore cannot engage the MLLE/PABC domain, was progressively displaced by poly(A) RNA. The independent PAM2–MLLE interaction is therefore sufficient to preserve PAIP1–PABPC1 binding when RNA occupies the RRM domains.

Since the nuclear poly(A)-binding protein PABPN1 also contains an RRM, we asked whether it is similarly displaced by PAIP1. Although recombinant PABPN1 bound A_18_ RNA with weaker affinity than PABPC1 (K_D_ ∼ 0.4 µM) (Figure S3J), PAIP1 did not displace PABPN1 from RNA (Figure S3K), consistent with the absence of a detectable PAIP1–PABPN1 interaction in Split-Venus experiments (Figure S3L).

Together, these findings support a model in which PAIP1 recruits PABPC1 to DDX19-containing export complexes. As PABPC1 starts to engage the poly(A)-tail of newly exported mRNPs, poly(A)+ RNA displaces the PAIP1 PAM1 motif from the PABPC1 RRM domains, while PAM2-MLLE binding retains PAIP1 in the complex. This bipartite interaction allows PAIP1 to remain associated with RNA-bound PABPC1 during cytoplasmic mRNP maturation.

### PAIP1 remodels the mRNP proteome and promotes PABPN1–PABPC1 exchange

If this model is correct, PAIP1 should promote association of PABPC1 with DDX19-containing export complexes. To test this prediction, we performed PABPC1–mCherry immunoprecipitation followed by mass spectrometry (IP-MS). As predicted, PAIP1 depletion substantially reduced co-purification of DDX19 and multiple nucleoporins (Figure 3A, S4A), indicating that PAIP1 connects PABPC1 to the mRNA export machinery.

**Figure 3.**
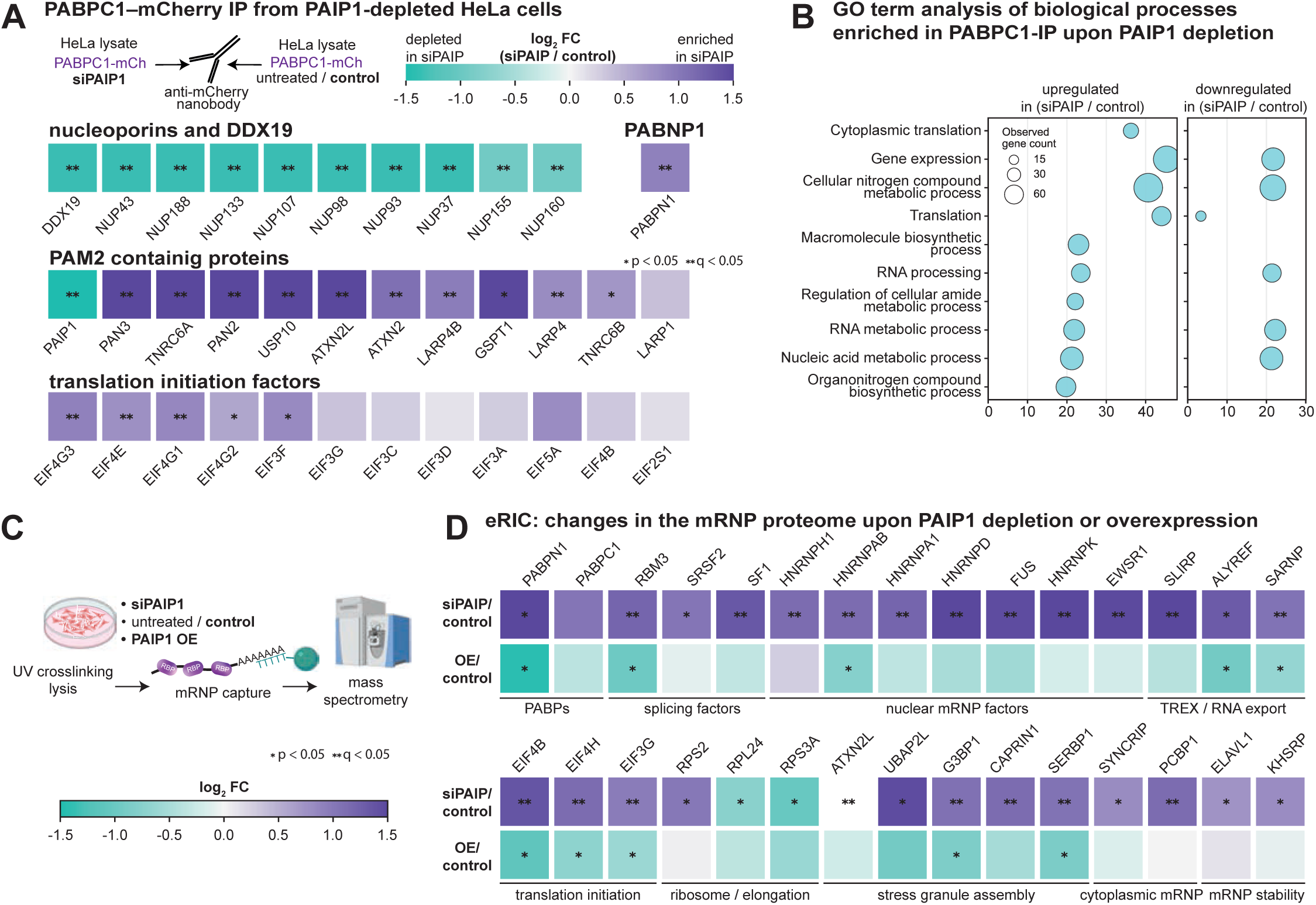
PAIP1 depletion modulates the PABPC1 interactome and cytoplasmic mRNP composition. (A) IP-MS of mCherry–PABPC1 interactomes in control and PAIP1-depleted cells, highlighting nucleoporins, PABPN1, PAM2-containing proteins and translation initiation factors. Extended views in Figure S4A. N = 3. (B) GO biological process enrichment analysis of differentially enriched proteins from Figure 3A. (C) Schematic of enhanced RNA interactome capture (eRIC). Following UV crosslinking and cell lysis, poly(A) RNAs and their associated proteins are captured using oligo(dT)-coupled beads. (D) Heatmap of proteins enriched or depleted in eRIC upon PAIP1 depletion or mild PAIP1 overexpression (∼1:1 relative to endogenous PAIP1 levels). Extended views in Figures S4B-F. N = 3.

Beyond these expected changes, PAIP1 depletion also substantially altered the broader PABPC1 interactome. Most notably, co-purification of PABPN1 with PABPC1 was strongly increased (Figures 3A and S4A). Because the immunoprecipitations were performed without nuclease treatment and no direct PABPN1–PABPC1 interaction has been reported, this likely reflects mRNPs carrying both PABPN1 and PABPC1. PAIP1 loss therefore increases mixed PABPN1/PABPC1 mRNPs, consistent with delayed or incomplete nuclear-to-cytoplasmic PABP exchange.

Association of PABPC1 with several PAM2-containing proteins (PAN3, ATXN2/L, and ERF3/GSPT1) also increased upon PAIP1 depletion, suggesting that PAIP1 normally occupies a substantial fraction of available PABPC1 MLLE domains (Figure 3A). Cytoplasmic translation was the most enriched GO biological process among the increased interactors, reflected by increased recovery of translation initiation factors (Figures 3A and 3B). Although PAIP1 has been proposed to bridge PABPC1 to EIF4G and EIF3^22–24^, their increased association with PABPC1 after PAIP1 depletion argues against a simple scaffolding role and instead suggests altered composition of PABPC1-containing mRNPs.

To determine whether PAIP1-dependent changes extend to poly(A)+ mRNPs, we performed enhanced RNA interactome capture (eRIC)^40^ in cells with endogenous PAIP1 levels, PAIP1 depletion, or mild PAIP1 overexpression (∼1:1 relative to endogenous protein) (Figure 3C). Following UV-crosslinking and preparation of whole-cell lysates (capturing both nuclear and cytoplasmic mRNPs), poly(A)+ RNA–protein complexes were isolated by oligo(dT) capture and analyzed by quantitative mass spectrometry. PABPN1 association with poly(A)+ RNA increased upon PAIP1 depletion (Figures 3D, S4B and S4F) and decreased upon PAIP1 overexpression (Figures 3D and S4D). By contrast, PABPC1 association was only mildly affected (Figures 3D, S4B and S4D), indicating that PABPN1 accumulated on poly(A)+ RNA without a corresponding loss of steady-state PABPC1 binding. PAIP1 therefore promotes removal of PABPN1 without being required for bulk steady-state PABPC loading. Since PAIP1 does not bind PABPN1, we propose that it promotes PABP exchange by bringing PABPC1 in close proximity to newly exported mRNPs, thereby facilitating displacement of PABPN1.

Beyond PABP exchange, PAIP1 abundance also broadly altered the poly(A)-associated proteome, with depletion and overexpression producing largely reciprocal effects (Figures 3D and S4B-F). PAIP1 depletion increased recovery of several nuclear RNA-processing proteins, including splicing and TREX/export factors. Because these factors are normally removed before or shortly after export, their increased recovery may reflect delayed mRNP maturation, and/or partial nuclear retention of mRNAs, despite no obvious bulk mRNA export defect (Figures 1L and 1M). PAIP1 depletion also increased recovery of cytoplasmic factors, including shuttling hnRNPs with known cytoplasmic functions, translation initiation factors, and stress-granule associated proteins, while ribosomal proteins were depleted. Together, these findings indicate that PAIP1 loss broadly alters the composition of poly(A)+ mRNPs beyond PABP exchange, potentially reflecting delayed establishment of a mature cytoplasmic mRNP state.

### PAIP1 levels tune multiple layers of mRNA metabolism

The broad changes in PABPC1- and poly(A)+ mRNP-associated proteomes prompted us to examine the consequences of PAIP1 depletion for mRNA abundance, half-life and poly(A)-tail length. We performed Oxford nanopore (ONT) cDNA sequencing to measure transcriptome-wide poly(A)-tail length, and SLAM-seq^41^ to quantify mRNA half-lives (Figure 4A and 4E). Steady-state mRNA abundance was obtained from the untreated SLAM-Seq input samples. All experiments were performed in control cells (48 h siCTRL treatment) and PAIP1-depleted cells (48 h siPAIP1 treatment).

**Figure 4.**
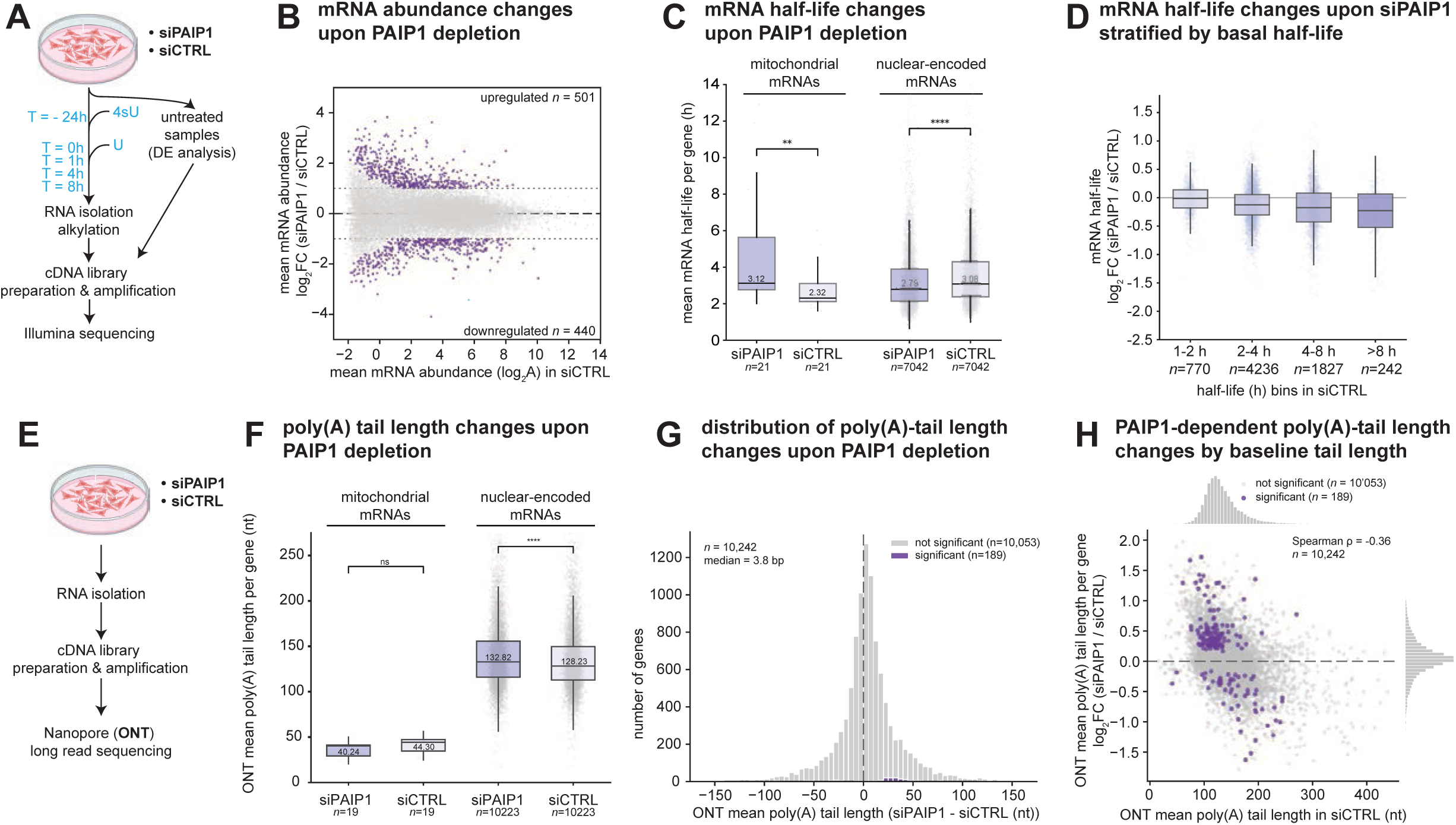
PAIP1 maintains poly(A)-tail homeostasis and mRNA stability genome-wide. (A) Schematic of the Illumina-based SLAM-seq mRNA half-life sequencing workflow. (B) MA plot of mRNA abundance log_2_ FC (siPAIP1/siCTRL) versus mean mRNA abundance (log_2_A) in siCTRL cells, measured by Illumina sequencing. Significantly changed genes are highlighted in violet. N = 3. (C) mRNA half-life estimates of mitochondrial and nuclear-encoded mRNAs in PAIP1-depleted (siPAIP1) or control-depleted (siCTRL) HeLa cells, measured by SLAM-seq. Significance was calculated using a Mann–Whitney test resulting in p-value 0.017 (*) or 6.6e−30 (****). N = 3. (D) Half-life FC (siPAIP1/siCTRL) for genes stratified by half-life in siCTRL cells. N = 3. (E) Schematic of the ONT-based poly(A)-tail length sequencing workflow. (F) Mean poly(A)-tail length of mitochondrial and nuclear-encoded mRNAs in PAIP1-depleted (siPAIP1) or control-depleted (siCTRL) HeLa cells, measured by ONT sequencing. Significance was calculated using a Mann–Whitney test resulting in p-value 0.32 (ns) or 2.4e^-36^ (****). N = 2. (G) Histogram of poly(A)-tail length changes upon PAIP1 depletion. Significantly changed genes are highlighted in violet. N = 2 (H) Correlation of mean poly(A)-tail length fold-change (FC; siPAIP1/siCTRL) plotted against mean poly(A)-tail length in siCTRL cells. Significantly changed genes are highlighted in violet. N = 2. (H) Distribution of poly(A)-tail length changes upon PAIP1 depletion. N = 2.

Differential expression (DE) analysis of steady-state mRNA abundance by edgeR identified *n* = 440 downregulated and *n* = 501 upregulated transcripts (at FDR ≤ 0.05, |log2FC| ≥ 1), including significant depletion of PAIP1 with log_2_FC = -3.4 (Figure 4B). Gene set enrichment analysis (GSEA) showed that genes downregulated upon PAIP1 depletion were enriched for spliceosome, ribosome biogenesis, and DNA replication functions (Figures S5A, −log_10_P ≥ 8), whereas upregulated transcripts were enriched for membrane, autophagy, and oxidative phosphorylation pathways, albeit at lower significance (−log_10_P ∼ 3-4) (Figures S5B).

Pulse-chase SLAM-seq (Figure 4A and Methods) yielded half-life estimates for *n* = 7’042 transcripts. Median transcript half-life decreased significantly from 3.08 hours in control cells to 2.79 hours in siPAIP1 cells (Figures 4C and S5C). Altered stability partially accounted for the observed steady-state abundance changes (Spearman correlation ρ = 0.32, n = 7’076 transcripts) (Figures S5D), indicating that PAIP1 loss modestly but broadly destabilizes cellular mRNAs. This effect depended strongly on baseline stability: short-lived transcripts with half-lives of 1-2 hours in wild type cells were least affected, whereas those with long half-lives (> 8 hours) were most strongly destabilized (Figures 4D).

Because PAIP1 promotes PABP exchange, we next examined poly(A)-tail length. Long-read nanopore sequencing (ONT) (Figure 4E) revealed a transcriptome-wide increase in median poly(A)-tail length from 127.54 nt in control cells to 132.32 nt after PAIP1 depletion (Figure 4F; Figures S5E-F for statistical controls). This effect was specific to nuclear-encoded mRNAs, as mitochondrial poly(A)-tail length remained largely unchanged (median = 41 - 43 nt, Figure 4F). At the level of individual transcripts, poly(A) length is frequently changed by several tens of nucleotides in either direction (Figure 4G). Although transcript-level significance was largely limited to well-covered transcripts (see methods), the genome-wide pattern was highly consistent: short poly(A) tails generally lengthened, whereas long tails tended to shorten (Figure 4H).

We next examined whether the magnitude of PAIP1-dependent responses could be explained by intrinsic transcript properties. For both mRNA half-life (Figure S5G) and poly(A)-tail length (Figure S5H), neither individual transcript features nor the combined multivariate models explained more than a small fraction of the observed variation in response magnitude. Since PAIP1 has previously been described as a translation coactivator, we examined translation efficiency in more detail. Cross-referencing our dataset with published HeLa translational-efficiency measurements^42,43^ revealed that the most efficiently translated transcripts exhibited the smallest PAIP1-dependent poly(A)-tail changes (Figure S5I), whereas mRNA half-life changes showed no comparable relationship with translation efficiency (Figure S5J). Consistent with this trend, ribosomal protein mRNAs, which are among the most efficiently translated transcripts, showed markedly smaller poly(A)-tail changes than all other mRNAs (Figure S5K). By contrast, re-analysis of published PABPC1/4-knockdown data^44^ showed the opposite relationship, with efficiently translated transcripts exhibiting the largest poly(A)-tail changes (Figure S5L). Thus, translation efficiency distinguishes the poly(A)-tail responses to PAIP1 and PABPC1/4 depletion.

Together, these analyses show that PAIP1 influences distinct layers of mRNA metabolism. Long-lived transcripts are preferentially destabilized upon PAIP1 depletion, whereas efficiently translated transcripts exhibit smaller poly(A)-tail changes. These findings indicate that PAIP1-dependent regulation of mRNA stability and poly(A)-tail length follows distinct transcript-specific patterns.

### Maternal PAIP1 is required for embryonic viability and proper patterning

The broad effects of PAIP1 depletion on mRNP composition, mRNA stability and poly(A)-tail length suggested that PAIP1 may be particularly important when gene expression depends predominantly on post-transcriptional regulation. Early embryogenesis provides such a context. Before zygotic transcription begins, development and embryonic patterning depend on maternally deposited mRNAs whose translation, poly(A)-tail length and stability are extensively regulated during the maternal-to-zygotic transition (MZT) and zygotic genome activation (ZGA) ^45–47^. Intriguingly, PAIP1 is highly expressed in germline tissues across species, including ovaries and oocytes of *Drosophila melanogaster*, *Danio rerio*, and *Homo sapiens*, and remains abundant during pre-ZGA development^48–53^ (Figure S6A).

This expression pattern suggests that maternally deposited PAIP1 may contribute to global post-transcriptional control, in particular poly(A)-tail homeostasis and translation, during early embryogenesis and the MZT. To test the physiological importance of maternal PAIP1, we depleted Paip1 specifically in the female germline of *Drosophila melanogaster* using matα-tub-Gal4-driven shRNA expression. After crossing these females to wild-type males, we analyzed the progeny at various stages of embryonic and post-embryonic development (Figure 5A). Whereas control embryos developed normally, maternal Paip1-KD caused pronounced embryonic lethality across three independent RNAi lines (TRiP.HM04067, GD13792, and KK107345), reaching lethality rates of 56%, 80%, and 74%, respectively (Figure 5B).

**Figure 5.**
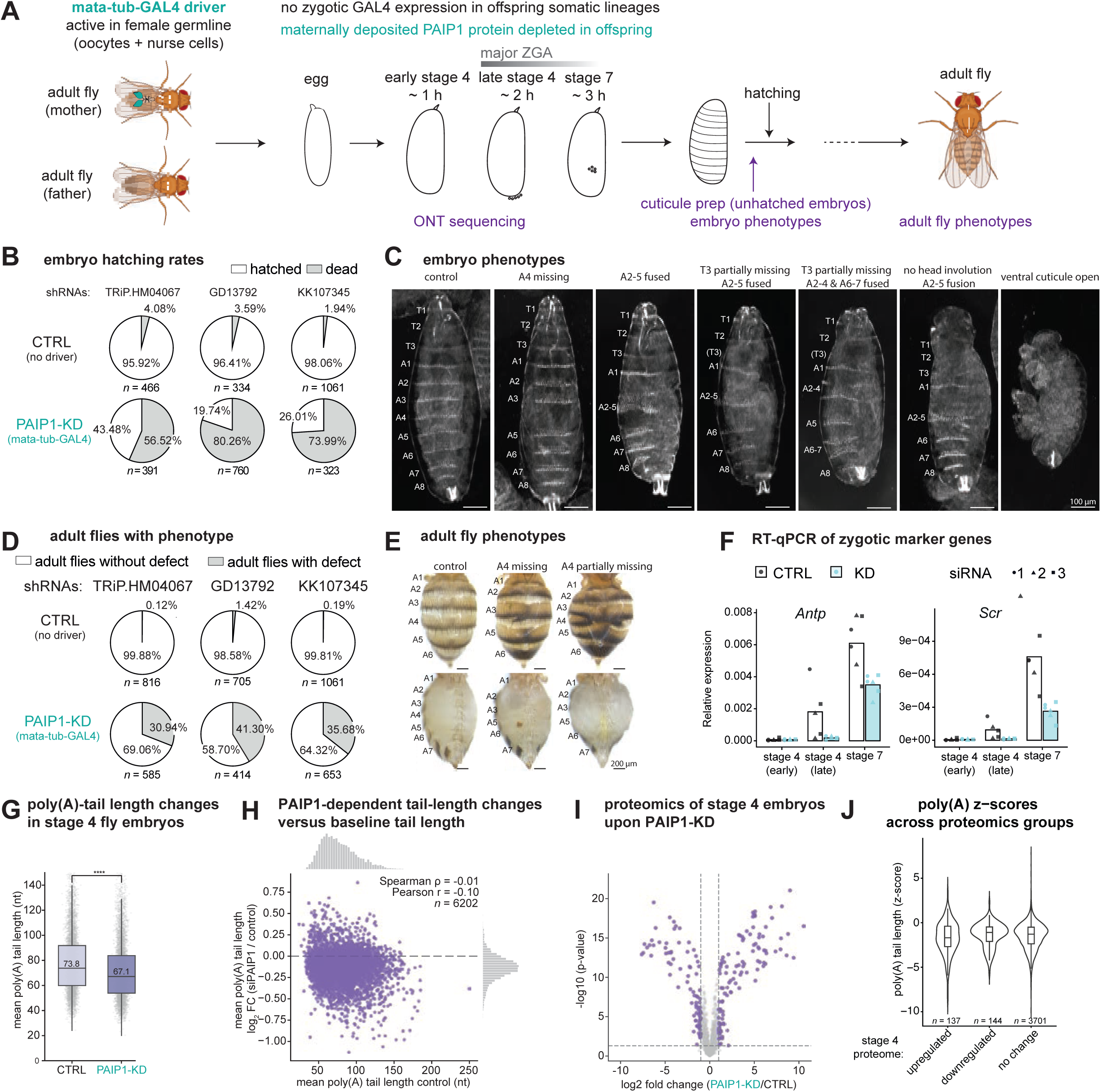
Maternal PAIP1 is required for *Drosophila* embryonic development and poly(A)-tail regulation. (A) Schematic of the maternal PAIP1 knockdown strategy using matα-tub-Gal4-driven shRNA expression in the female germline, with the timing of experimental analyses during embryonic development indicated. (B) Embryo hatching rates for three independent shRNA lines expressed in the maternal germline, comparing control (CTRL, lacking driver) and PAIP1-knockdown (PAIP1-KD) embryos. N = 3. (C) Representative dark-field images of cuticular preparations from embryos hatched from CTRL or PAIP1-KD embryos. Scale bars, 100 µm. (D) Percentage of adult flies displaying developmental abnormalities, derived from CTRL or PAIP1-KD embryos as in (B). N = 3. (E) Representative images of adult flies from. Scale bars, 200 µm. (F) RT-qPCR of zygotic marker genes at embryonic stages early 4, late 4, and 7 in CTRL and PAIP1-KD embryos. Data represent mean ± SD; N = 3. (G) Mean mRNA poly(A)-tail length in stage 4 embryos of CTRL and PAIP1-KD mothers, determined by ONT sequencing. N = 3. (H) Scatter plot of poly(A)-tail length FC (PAIP1-KD/CTRL) versus poly(A)-tail length in CTRL embryos. N = 2. (I) Scatter plots of protein abundance FC (PAIP1-KD/CTRL) at embryonic stage 4 versus at stage 7, with color indicating poly(A)-tail length log_2_ FC (PAIP1-KD/CTRL) (I). N = 6. (J) poly(A)-tail length changes (Z-scores) for genes encoding proteins that are upregulated, downregulated, or unchanged in stage 4 proteomics data. N = 6.

Cuticle preparations, which report the terminal phenotype of unhatched embryos, revealed a spectrum of anterior–posterior patterning defects, including loss or disruption of individual abdominal segments, most frequently the A4 denticle belt, and partial or complete fusions within the A2–A5 region (Figure 5C). More severe phenotypes displayed posterior segment fusions, e.g. A6–A7 and A4-5, together with anterior morphogenesis defects such as lack of thoracic segment T3 or failed head involution (Figure 5C). The PAIP1-KD embryos that completed embryogenesis and hatched retained related abdominal segmentation abnormalities at the adult stage (Figures 5D and 5E), indicating that patterning defects established early during embryogenesis persist into later developmental stages. Maternal PAIP1 is therefore broadly required for anterior–posterior patterning.

These phenotypes were preceded by altered early embryonic gene expression. RT-qPCR showed that maternal Paip1-KD perturbed induction of embryonic patterning genes (*Antp*, *eve*, *salm*, and *Scr*) as well as the purely zygotic transcript *roX1* (Figures 5F and S6B). The defects were already apparent early in development (stage 4), indicating delayed establishment of zygotic gene expression programs. This is particularly striking because zygotic Paip1 mutants, in which maternally deposited Paip1 remains present during early embryogenesis, progress beyond the MZT and arrest only days later at the pupal stage^54^. Together with the maternal and therefore transient nature of our knockdown, this distinction identifies an early requirement for maternal PAIP1 during the MZT.

### Loss of maternal PAIP1 shortens poly(A)-tails

The phenotypic spectrum of maternal PAIP1 depletion resembles defects reported after perturbation of Wispy (*wisp*), the *Drosophila* cytoplasmic poly(A) polymerase, which likewise causes segmental deletions and fusions through altered poly(A)-tail regulation of maternal transcripts, including *bicoid*^55,56^. This similarity raised the possibility that maternal PAIP1 contributes to poly(A)-tail regulation during early embryogenesis.

We therefore measured poly(A)-tail lengths transcriptome-wide by nanopore cDNA sequencing of RNA isolated from stage 4 embryos, before zygotic transcription is fully established. Among 6’210 detected transcripts, 1’223 (20 %) showed significant poly(A)-tail shortening upon maternal Paip1 knockdown, whereas only 19 (0.3 %) exhibited tail extension (Figures 5G and 5H). These changes occurred without corresponding alterations in bulk RNA abundance, which remained largely unchanged, as expected at this predominantly transcriptionally inactive pre-ZGA stage (Spearman corr = 0.02; Figures S6C and S6D). Interestingly, the relationship between translational efficiency^57^ and PAIP1-dependent tail changes was also observed in embryos and even more pronounced than in our HeLa dataset: poorly translated maternal transcripts showed significantly greater tail changes than efficiently translated transcripts (Spearman ρ = −0.14 for 0–1 h embryo; Spearman ρ = −0.05 for 2-3 h embryo) (Figure S6E).

Because poly(A)-tail length is a major determinant of maternal mRNA translation during early embryogenesis^47,57^, we next analyzed protein abundance in stage 4 and stage 7 embryos. Maternal PAIP1 depletion caused broad proteome changes at both stages (88 and 109 increased, 190 and 83 decreased proteins at stages 4 and 7 respectively, |log_2_FC| ≥ 1 and p-value < 0.05) (Figures 5I, S6F and S6G). Unexpectedly, upregulated proteins tended to be encoded by transcripts with stronger poly(A)-tail shortening than those encoding unchanged or downregulated proteins (Figure 5J). This contrasts with the positive coupling between tail length and translation generally observed in early embryos^44,57,58^, but resembles the high translational efficiency of short-tailed transcripts reported in somatic cells^59^. These observations point to poly(A)-tail length and protein abundance becoming at least partially uncoupled upon maternal PAIP1 depletion.

A focused analysis of anterior–posterior patterning factors showed that PAIP1 depletion altered the poly(A)-tail length of several early maternal regulators, including *smg*, *png*, *CycB* and *swa* (Figure S6H). Correspondingly, protein abundance of maternal patterning proteins, including Bicoid, Caudal, Hunchback, Swallow and Torso, was altered by stage 7 (Figure S6I). Because the poly(A)-tail analysis was performed before ZGA, most downstream zygotic patterning factors were absent or only weakly detected and could not be assessed.

Together, these findings demonstrate that maternal PAIP1 maintains poly(A)-tail homeostasis across a substantial subset of maternal transcripts. Its loss delays activation of zygotic developmental programs, perturbs upstream patterning determinants and alters proteome composition, consistent with defective establishment of translation-competent maternal mRNPs.

## DISCUSSION

Export through the NPC marks the transition between nuclear and cytoplasmic stages of gene expression, yet how newly exported mRNPs acquire their cytoplasmic protein composition has remained one of the least understood transitions in RNA biology. Our findings identify PAIP1 as a DDX19-associated adaptor that couples the final steps of mRNA export to cytoplasmic mRNP maturation.

DDX19 emerges as a pore-proximal platform for cytoplasmic mRNP maturation alongside its established role in mRNA export. The N-terminal SBM motif of DDX19 provides a conserved recruitment platform for MIF4G-domain cofactors, including PAIP1, MIF4GD/SLIP1, and CTIF, thereby serving as a molecular hub that couples mRNA export to distinct cytoplasmic processes, including poly(A)-dependent mRNP maturation and translation, histone mRNA metabolism, and pioneer translation, respectively^20,21,60,61^ (Figure S7A). The importance of this interface is underscored by viral co-option: the SARS-CoV-2 protein NSP3 binds the PAIP1 MIF4G domain through the same SBM-binding groove^31^ (Figure S1F). Together with recent work showing that SARS-CoV-2 recruits PABPC1 to stabilize its own mRNA^62^, this raises the possibility that coronaviruses exploit the PAIP1-PABPC1 module to favor assembly of translation-competent viral mRNPs.

How might PAIP1 promote the PABPN1-to-PABPC1 exchange? We did not detect a direct PAIP1–PABPN1 interaction, arguing against an active removal mechanism. Exchange is likely to be thermodynamically favorable once transcripts reach the cytoplasm, where PABPC1 is more abundant and binds poly(A) RNA with higher affinity than PABPN1. The challenge may therefore be kinetic: not whether exchange occurs, but how rapidly newly exported mRNPs acquire PABPC1. We propose that PAIP1, possibly together with DDX19-mediated PABPN1 remodeling^13^, accelerates this process by positioning PABPC1 close to the nuclear pore. Self-association of both PAIP1 (our data) and PABPC1^63^ may further promote cooperative loading of multiple PABPC1 molecules, progressively displacing residual PABPN1 and stabilizing the cytoplasmic mRNP state (Figure S7B). Such a gradual transition would be consistent with recent time-resolved eRIC analyses, which indicate that the PABPN1-to-PABPC1 transition proceeds progressively during the early cytoplasmic phase rather than occurring immediately upon export^64^.

This model also reframes PAIP1’s established role in translation. PAIP1 has long been viewed as a translation-activating PABPC1 cofactor^22–24^, but our data suggest that its function begins earlier, by establishing a translation-competent cytoplasmic mRNP after export. Translation activation may therefore represent a downstream consequence of an earlier export-coupled maturation step that establishes the cytoplasmic identity of mRNPs and shapes their subsequent regulation. The presence of an SBM motif in EIF3G^20^ further suggests that PAIP1 may be recruited to the translation-initiation machinery through the same interaction principle that initially positions it at the nuclear pore.

Loss of PAIP1 has broad consequences for post-transcriptional gene regulation, impairing PABP exchange and the establishment of a mature cytoplasmic mRNP state, with consequences for both poly(A)-tail homeostasis and mRNA stability. Long-lived transcripts are preferentially destabilized, whereas the magnitude of poly(A)-tail changes decreases with increasing translation efficiency. This suggests that PAIP1-dependent defects in cytoplasmic mRNP maturation are manifested differently at the level of poly(A)-tail homeostasis and mRNA stability.

PAIP1-dependent mRNP maturation becomes particularly important when gene expression is controlled predominantly after transcription. Early embryogenesis provides such a context: before zygotic genome activation, development relies on maternally deposited mRNAs whose translation and stability are regulated through coordinated changes in poly(A)-tail length and mRNP composition^45,47^. Maternal PAIP1 depletion causes widespread poly(A)-tail shortening, delayed activation of zygotic gene-expression programs, and severe developmental patterning defects. Because PAIP1 is depleted maternally in our experiments, these defects are likely established during oogenesis, when maternal mRNAs are synthesized and exported. Early embryos may therefore be particularly vulnerable to defects in the establishment of maternal mRNP states, as their dependence on a finite maternal mRNA pool and limiting PABPC availability^44^ may increase the sensitivity of the embryo to defects in PAIP1-dependent mRNP maturation.

Together, our findings identify PAIP1 as a molecular link between mRNA export, PABP exchange, and cytoplasmic mRNP maturation. They further suggest that export-coupled mRNP maturation and translation activation are not distinct activities, but successive stages of the same process by which newly exported mRNAs acquire their cytoplasmic ‘identity’. Rather than a passive consequence of NPC passage, the nuclear-to-cytoplasmic transition emerges as an actively coordinated step that shapes downstream mRNA metabolism. The conservation of the PAIP1–DDX19 interface across metazoans suggests that this coupling represents a fundamental layer of post-transcriptional gene regulation.

## RESOURCE AVAILABILITY

Lead contact: Maria Hondele

### Materials availability

Plasmids, oligonucleotides and cell lines are available through the lead contact upon request.

### Data availability

Imaging data will be deposited in the BioImage Archive (EMBL-EBI). Raw sequencing data, including Oxford Nanopore are deposited in the GEO database and at the European Nucleotide Archive (ENA). Proteomics data are deposited at MassIVE. Accession numbers will be provided upon publication.

### Code availability

Code for the analysis and configuration of “nanoflowz” is available at https://github.com/zavolanlab/ONT_seq_PAIP1_Hondele release v1.0.0. Corresponding Zenodo archive of the release can be found here: https://doi.org/10.5281/zenodo.20729199.

## ACKNOWLEDGEMENTS

We thank Anne Spang, Susan E. Mango, Franka Voigt, Elisa Dultz, Peter Scheiffele and all members of the Hondele laboratory for insightful discussions and critical reading of the manuscript. We are grateful to Janani Durairaj, Stefan Bienert and Gerardo Tauriello for continuous support with AlphaFold predictions, and to Rebecca Talev, Arturo Sanchez and Niko Dietz for assistance with initial experiments. We thank the Bioinformatics, Biophysics (especially Tim Sharpe and Tobias Mühlethaler), Proteomics, Imaging Core Facility (IMCF) and sciCORE facilities at the University of Basel for their outstanding support throughout this project. We also thank the Functional Genomics Center Zurich (FGCZ), especially Anna Bratus, for their excellent support with ONT sequencing experiments. Computational analyses were performed at the sciCORE scientific computing center at the University of Basel. We thank Annemarie Overwijn for assistance with the graphical model. DO, MJG and MZe were supported by Fellowships from the International PhD Program in Molecular Life Sciences, Biozentrum, University of Basel. ITC measurements were supported by the SNSF R’Equip Grant 213436, “Efficient Characterization of Biomolecular Interactions by Automated Isothermal Titration Calorimetry.” MH acknowledges funding from the Swiss National Science Foundation (PCEFP3_187052) and the European Research Council (ERC Starting Grant 950262). MH, CIKV, MZa and TM received institutional funds from the Kanton Basel-Stadt and Basel-Land provided to the Biozentrum of the University Basel.

## AUTHOR CONTRIBUTIONS

conceptualization, D.O., C.I.K.V., and M.H.;

methodology, D.O, A.M, R.I., P.S., M.Ze. and M.S.

formal analysis, D.O., K.D, M.M., A.M, R.I., M.J.G., C.I.K.V. and M.H.;

investigation, D.O., K.D, C. d. A., M.M, A.M, R.I. M.J.G, I.S. and S.B.;

resources, K.D, M.M., I.S. and N.B.;

writing - original draft, D.O., C.I.K.V. and M.H.;

writing - reviewing & editing, K.D., C. d. A., A.M, M.J.G, and M. Za;

visualization, D.O, K.D, M.M, C.I.K.V. and M.H.;

supervision, T.M., M.Za., C.I.K.V., and M.H;

funding acquisition, T.M., M.Za., C.I.K.V., and M.H.

## DECLARATION OF AI AND AI-A

During the preparation of this work, the author(s) used ChatGPT and CLAUDE.ai to improve wording and the clarity of the manuscript and to assist with script writing. The author(s) reviewed and edited the content as appropriate and take full responsibility for the content of the publication.

## SUPPLEMENTAL INFORMATION

Document S1: Figures S1-S6

Supplemental Table 1: relative abundance of PAIP1 and components of the DDX19–PABP exchange pathway in different cell lines and tissues

Supplemental Table 2: oligonucleotides used in this study

## SUPPLEMENTARY FIGURE LEGENDS

**Figure S1. Structural and evolutionary analysis of the PAIP1–DDX19 interface.** (A) AlphaFold-Multimer screen of DDX19A/B against their BioGRID interactome, ranked by the iPTM or 0.8 iPTM + 0.2 PTM score, using 0.8 and 0.7 as reference for reliable predictions, respectively. (B) AlphaFold-Multimer screen of PABPC1 against the 30 BIOGRID interactors shared between PABPC1 and DDX19 (see Figure 1A), ranked by the iPTM, pdockq2 or 0.8 iPTM + 0.2 PTM score. (C) Domain architecture of DDX19, the MIF4G proteins PAIP1, GLE1, CTIF and MIF4GD/SLIP1, and the poly(A) binding proteins PABPN1 and PABPC1. (D) Overlay of multiple AlphaFold-Multimer models of DDX19B in complex with the PAIP1 MIF4G domain and corresponding PAE plots. (E) Full-length PAIP1 is predicted by AlphaFold-Multimer to bind DDX19A and DDX19B through a similar interface as the isolated PAIP1 MIF4G domain, as shown by structural models and corresponding PAE plots. (F) Overlay of an AlphaFold model of the DDX19 SBM motif in complex with the PAIP1 MIF4G domain, and the crystal structure of NSP3 bound to the PAIP1 MIF4G domain (PDB: 6YXJ). (G) Presence of DDX19, GLE1, PAIP1, CTIF, and MIF4GD/SLIP1 across representative eukaryotic lineages. (H) Phylogenetic tree of PAIP1. (I) AlphaFold models of PAIP1–DDX19 complexes from early-branching metazoans and corresponding PAE plots. (J) Isothermal titration calorimetry (ITC) measurements for PAIP1, MIF4GD/SLIP1 or CTIF titrated into DDX19B; corresponding to summary table in Figure 1F. (K) Volcano plot of immunoprecipitation - mass spectrometry (IP-MS) of stable HeLa cell lines expressing V5–DDX19B or V5–DDX19B^Δ2–12^ at endogenous level; SBM-binding proteins are highlighted. N = 3.

**Figure S2. Biochemical validation of the PAIP1–DDX19 interaction.** (A) In vitro ATPase activity of recombinant DDX19A/B in the presence or absence of poly(A) RNA and recombinant PAIP1 as indicated, significance was determined using a one-way ANOVA with Tukey’s multiple-comparisons test. N = 3. (B) Immunoblot showing expression of the siRNA-resistant PAIP1– mEGFP constructs used in Figures 1I and 2D and at levels comparable to endogenous PAIP1. N = 3. (C) Schematic of the Split-Venus bimolecular fluorescence complementation (BiFC) assay. (D,E) Split-Venus bimolecular fluorescence complementation (BiFC) analysis in HeLa cells transiently co-transfected with DDX19A-VC (D) or DDX19B-VC (E) with GLE1-VN, CTIF-VN, and MIF4GD/SLIP1-VN; cells were co-stained for LaminA/C. Scale bar: 10 µm, N = 3, n ≥ 36 cells. (F) Poly(A) RNA FISH in HeLa cells treated with control siRNA or siRNA targeting DDX19A/B, with or without expression of DDX19^WT^ or DDX19^Δ2–12^ constructs. Scale bars 10 µm, N = 3, n ≥ 100 cells. (G) Quantification of nuclear-to-cytoplasmic poly(A) RNA signal (median intensity ratio) from cells shown in S2F. Statistical comparisons were performed on replicate-level median nuclear-to-cytoplasmic poly(A) intensity ratios using a linear model with biological replicate as a blocking factor, followed by Dunnett-adjusted comparisons against siCNTL. Only siDDX19 differed significantly from control (p=2.53×10^−6^); all other comparisons were not significant. (H) Immunoblot showing knockdown of endogenous DDX19A/B and expression of siRNA-resistant DDX19–V5 constructs used in Figure S2F, at levels comparable to endogenous DDX19. N = 3. (I) Immunoblot showing knockdown of endogenous PAIP1 used in Figure 1L. N = 3.

**Figure S3. Structural and functional characterization of the PAIP1–PABPC1 interaction.** (A) Schematic representation of domain architecture and interaction interfaces of PAIP1 and PABPC1, with PAIP1^mutPABPC^ mutations indicated; AlphaFold-Multimer model of full-length PAIP1 bound to PABPC1, with point mutations highlighted; (B) Structural overlay of the PAIP1–PABPC1 RRM2 interface with the crystal structure of RNA-bound PABPC1 (PDB: 4F02)^65^. (C) Structural overlay of the PAIP1-PABPC1 interface with the crystal structure of RNA-bound PABPC1 RRM2 (PDB: 4F02), superimposed on RRM3. (D) Comparison of the predicted PAIP1 PAM2–MLLE interface with the previously reported NMRstructure of the complex (PDB: 1JH4)^38^. (E) Alphafold prediction of two full-length PAIP1 molecules; insert shows dimerization interface. (F) SEC-MALS of the PAIP1 MIF4G domain indicates dimerization. (G) HeLa cell lines stably expressing PAIP1^WT^–mEGFP or PAIP1^mutPABPC^-mEGFP were subjected to anti-GFP immunoprecipitation followed by mass spectrometry to identify enriched proteins; nucleoporins and PABPC proteins are highlighted. N = 3. (H) Fluorescence polarization analysis of PABPC1 binding to A18 RNA. Concentration marked with an arrow (12 nM) was used for assays in Figure 2G. N = 3. (I) Fluorescence polarization analysis of PABPN1 binding to A18 RNA. Concentration marked with an arrow (1 µM) was used for assays in Figure S3J. N=3. (J) Fluorescence polarization competition assay in which recombinant PAIP1 were titrated against a fixed concentration of a preformed A18 RNA–PABPN1 complex (see Figure S3G), or RNA alone. N=3. (K) Split-Venus analysis of cells transiently transfected with PAIP1-VC and PABPC1-VN, or PAIP1-VN with PABPN1-VC. Scale bars 10 µm, N=3, n ≥ 26 cells.

**Figure S4. eRIC proteome analysis of PAIP1-depleted and PAIP1-overexpressing cells.** (A) IP-MS of mCherry–PABPC1 interactomes in control and PAIP1-depleted cells, highlighting nucleoporins (left), PAM2-containing proteins (middle) and PABPN1 and translation initiation factors (right). Same data as heatmap in Figure 3A. N = 3. (B) eRIC proteome of PAIP1-depleted versus control cells. Same data as Figure 3D. N = 3. (C) GO cellular component enrichment analysis of Figure 3A and S4B. (D) eRIC proteome of 1:1 PAIP1 overexpressing versus control cells. Same data as Figure 3D. N = 3. (E) GO cellular component enrichment analysis of Figure 3A and S4D. (F) Extended heatmap of eRIC enrichment. Same data as Figure 3D.

**Figure S5. Additional analyses of mRNA stability, poly(A)-tail length, and transcript abundance following PAIP1 perturbation.** (A) GSEA of genes downregulated upon PAIP1 depletion, from Illumina RNA-seq data (Figure S6A). N = 3. (B) GSEA of genes upregulated upon PAIP1 depletion, from Illumina RNA-seq data (Figure S6C). N = 3. (C) Scatter plots comparing mRNA half-life estimates in siPAIP1 and siCTRL cells. Significantly changed transcripts are highlighted in violet. N = 3. (D) Scatter plots comparing mRNA half-life estimates and differential expression in siPAIP1 and siCTRL cells. genes show a partial correlation, ⍴ = 0.32, n = 7076. N = 3. (E) Biological replicate consistency of ONT-derived poly(A)-tail measurements. Mean per-gene poly(A)-tail lengths were compared between biological replicates for siCTRL (left) and siPAIP1 (right). Analysis was restricted to genes supported by at least 10 ONT reads in each sample (n = 5,685 genes). Pearson correlation coefficients are indicated. (F) Randomization test for transcriptome-wide poly(A)-tail lengthening following PAIP1 depletion. Condition labels were randomly permuted among the four biological replicates (two siCTRL, two siPAIP1), and the transcriptome-wide median poly(A)-tail length difference (siPAIP1 − siCTRL) was recalculated for 2,000 permutations to generate the null distribution (grey histogram). The observed median difference (+2.84 nt; purple line) falls outside the null distribution (p < 0.0005), supporting a robust global increase in poly(A)-tail length upon PAIP1 depletion. (G,H) Association of transcript features with PAIP1-dependent changes in mRNA half-life (G) and poly(A)-tail length (H). Left, Spearman correlation coefficients (ρ) between individual transcript features and the magnitude of mRNA half-life or poly(A)-tail length changes upon PAIP1 depletion (|log₂ fold change|, siPAIP1/siCTRL). Right, all transcript features were jointly entered into multivariable rank regression models predicting the magnitude of each response variable. Points represent standardized regression coefficients (β); error bars indicate bootstrap 95% confidence intervals. Feature definitions and data sources are described in the Methods. (I,J) Association between translational efficiency and PAIP1-dependent changes in poly(A)-tail length (I) and mRNA half-life (J). Transcripts were grouped into quintiles according to published translational efficiency measurements^43^. Boxplots show the distribution of the magnitude of poly(A)-tail length changes (I) or mRNA half-life changes (J) upon PAIP1 depletion (|z-score(siPAIP1/siCTRL)|, noise-normalized). Dots represent individual transcripts. Spearman correlation coefficients (ρ) and the number of transcripts analyzed (n) are indicated. (K) Comparison of PAIP1-dependent poly(A)-tail length log₂ fold-changes between ribosomal-protein mRNAs and all other transcripts. P value 1.3x10-17, Mann–Whitney U test. (L) Median absolute poly(A)-tail length change (nucleotides) following PABPC1/4 depletion across translational-efficiency quintiles in HeLa cells. Published PABPC1/4 depletion data^44^ were reanalyzed using the same transcript stratification as in Figures S5I and S5J.

**Figure S6. Extended analysis of poly(A)-tail regulation and zygotic gene expression in PAIP1-KD *Drosophila* embryos.** (A) PAIP1 mRNA expression in fly, zebrafish and human germline and early development. Data from^48–50,52^. (B) RT-qPCR of zygotic marker genes at embryonic stages early 4, late 4, and 7 in CTRL and PAIP1-KD embryos. See also Figure 5F. Data represent mean ± SD; N = 3. (C) MA plot of mRNA abundance log_2_ FC (PAIP1-KD/CTRL) versus mean mRNA abundance in CTRL embryos. N = 3. (D) Scatter plot of poly(A)-tail length log_2_ FC (PAIP1-KD/CTRL) versus mRNA abundance log_2_ FC (PAIP1-KD/CTRL) at embryonic stage 4. N = 3. (E) Comparison of PAIP1-dependent poly(A)-tail length log₂ fold-change with translational efficiency stratified into quintiles. Translational efficiency was obtained from published *Drosophila* embryo ribosome profiling data^57^. (F) Volcano plots of protein abundance log₂ fold-change (PAIP1-KD/CTRL) at embryonic stage 7. N = 3. (G) Scatter plots of protein abundance log₂ fold-change (PAIP1-KD/CTRL) at embryonic stage 4 versus at stage 7. N = 3. (H,I) PAIP1-dependent poly(A)-tail (H) and proteome (I) changes across the *Drosophila* anterior– posterior segmentation network. The canonical segmentation network is shown as directed graph comprising four regulatory tiers: maternal (*CycB*, *bcd*, *cad*, *exu*, *nanos*, *png*, *smg*, *swa*, and *tor*), gap (*Kr*, *gt*, *hb*, *hkb*, *kni*, and *tll*), pair-rule (*eve*, *ftz*, *hairy*, *odd*, *opa*, *prd*, *run*, and *slp1*), and segment-polarity (*en* and *wg*) genes. Edges represent established regulatory interactions from developmental genetics (18 activating, blue; 29 repressing, orange) and are shown as a reference network rather than inferred from these data. Node color indicates the log₂ fold change in PAIP1 knockdown relative to control. (H) Poly(A)-tail length changes in stage 4 embryos. (I) Protein abundance changes in stage 7 embryos. Bold outlines indicate significant changes (poly(A): adjusted P < 0.05; protein: q < 0.01), thin outlines indicate quantified but non-significant genes, and grey hatched nodes indicate genes that were not detected in the respective dataset. Detection and statistical significance are panel-specific.

**Figure S7. Model for PAIP1-mediated coupling of mRNA export to cytoplasmic mRNP maturation.** (A) DDX19 acts as a pore-proximal docking platform for MIF4G-domain cofactors through its N-terminal SBM motif. While CTIF and MIF4GD/SLIP1 recruit DDX19 to specialized translation pathways, PAIP1 links DDX19 to the canonical poly(A)-dependent translation machinery through its interaction with PABPC1. (B) Proposed model for PAIP1-mediated PABP exchange. PAIP1 is recruited to DDX19 at the cytoplasmic face of the nuclear pore through the SBM motif and delivers PABPC1 to newly exported mRNPs. As PABPC1 engages the poly(A)-tail, residual PABPN1 is progressively displaced, while PAIP1 remains associated with PABPC1 through its PAM2–MLLE interaction. PAIP1 and PABPC1 self-association may further promote cooperative loading of additional PABPC molecules, thereby facilitating establishment of a mature cytoplasmic mRNP state.

## Key resources table

see Supplemental Table 2

## Method details

### Alphafold 2 Screens

Protein–protein interaction screening was performed using Alphafold multimer v2.3 ^1^, in which multiple sequence alignments are generated on CPUs and structure predictions are carried out on GPUs. For each of the DDX19 linked 251 candidate interactors obtained from BioGRID v5.0, five structural models were predicted in complexes with either DDX19A or DDX19B. Interaction confidence metrics, such as pDOCKq2 2, 0.8 iPTM + 0.2 PTM and iPTM were computed and extracted, after which they were plotted using jupyter notebooks. Predictions with an iPTM score ≥ 0.8 or a pDockQ2 score ≥ 0.23 and iPTM+PTM ≥0.7 were considered reliable models. Corresponding structural models and predicted aligned error (PAE) plots were subsequently inspected manually. Structural models and PAE plots were visualized using UCSF ChimeraX (v1.11).

### Alphafold 3 modeling

Protein–protein complex structures were predicted using AlphaFold 3 (as annotated in figure legends) using the AlphaFold Server web interface (alphafoldserver.com^3^). Amino acid sequences of the interacting proteins were submitted as separate protein chains using the default server settings. The resulting structural models and predicted aligned error (PAE) plots were inspected and visualized using UCSF ChimeraX (v1.11). The models and PAE plots were displayed using UCSF ChimeraX (v1.11).

### Plasmid construction

All proteins for expression in human cells were cloned into a pcDNA5 vector, with their respective tags. Constructs for recombinant expression in *E. coli* and purification were cloned into pETMCN vectors containing their respective tags.

### Protein expression and purification

Recombinant protein expression was performed in E. coli Lemo21(DE3) cells. Bacteria were transformed and grown overnight at 37 °C in precultures containing the appropriate antibiotics. Expression cultures were then inoculated and grown at 37 °C to an OD_600_ of 0.5 – 0.9, induced with 200 µM IPTG, and incubated overnight at 19 °C prior to harvest. Bacterial pellets were lysed by pressure homogenization using an EmulsiFlex system (Avestin C5) in lysis buffer containing 25 mM potassium phosphate (pH 7.5), 500 mM NaCl, 10% (w/v) glycerol, protease inhibitors, DNaseI, and RNaseA. Lysates were clarified by centrifugation (70,000 × g, 30 min, 4 °C) followed by filtration through a 0.45 µm filter after which imidazole was added to a final concentration of 20 mM. Proteins were first affinity purified by immobilized metal affinity chromatography (IMAC) using a homemade 5-mL column packed with Ni Sepharose High Performance resin (Cytiva). The clarified lysate was loaded onto the column, which was subsequently washed with at least 10 column volumes of wash buffer (25 mM potassium phosphate, pH 7.5, 500 mM NaCl, 10% (w/v) glycerol, and 50 mM imidazole). Bound proteins were eluted using a linear imidazole gradient to a final concentration of 500 mM.Proteins used for pulldown experiments were directly subjected to size-exclusion chromatography (SEC) using a 16/600 HiLoad Superdex 200 pg column (Cytiva) equilibrated in SEC buffer (25 mM potassium phosphate pH 7.5, 500 mM NaCl, 10% (w/v) glycerol, 0.5 mM β-mercaptoethanol). From all other proteins, the purification tags were removed by HRV-3C protease cleavage either overnight at 4 °C or for 3 h at room temperature in SEC buffer, or, in the case of PABPN1 and PABPC1, in dialysis buffer (25 mM potassium phosphate pH 7.5, 150 mM NaCl, 10% (w/v) glycerol, 0.5 mM β-mercaptoethanol). Following cleavage, uncleaved proteins, protease, and residual affinity tags were removed by reverse IMAC. Cleaved PABPC1 and PABPN1 proteins were subsequently diluted to a final salt concentration of 75 mM NaCl and further purified by ion-exchange chromatography using a MonoQ 10/100 GL column, followed by SEC as described above. All other proteins were directly purified by SEC using 16/600 HiLoad Superdex 200 or Superdex 75 pg columns (Cytiva), as described above.

SEC fractions were analyzed by SDS page; clean fractions were pooled and concentrated using Amicon ultra centrifugal filters with a cutoff of half or less the molecular weight of the protein.

### Pull-down Assays using recombinant proteins

Recombinant MBP-tagged proteins were mixed with equimolar amounts (10 or 20 µM) of binding partners (= input) in wash buffer (PBS supplemented with 1 mM MgCl₂ and 0.1% Tween-20) and incubated with pre-equilibrated amylose resin at 4 °C for 1–2 h. The resin was collected by centrifugation (800 xg, 3 min) and washed three times with wash buffer. Bound proteins were eluted by 15 min incubation with elution buffer (PBS supplemented with 1 mM MgCl₂, 0.1% Tween-20, and 10 mM maltose), followed by centrifugation to separate the beads, and the supernatant was collected (= elution). Input and elution fractions were analyzed by SDS–PAGE and visualized by Coomassie staining.

### Pull-down competition assays in the presence of RNA

MBP pulldown assays in the presence of RNA were performed using titrated poly(A) RNA at concentrations of 0.001, 0.01, 0.1, or 1 µg µL⁻¹, with RNasin added according to the manufacturer’s recommendations (Promega). All other steps were performed as described for the standard pulldown assays.

### Isothermal Calorimetry

Proteins used for isothermal titration calorimetry (ITC) were dialyzed into binding buffer consisting of 25 mM potassium phosphate (pH 7.5), 150 mM NaCl, 1 mM MgCl₂, 0.5 mM TCEP, and 10% (w/v) glycerol. ITC measurements were performed using a MicroCal PEAQ-ITC Automated instrument (Malvern Panalytical) at 25 °C. DDX19B at a concentration of 20 µM was loaded into the sample cell, while binding partners (the MIF4G domains of PAIP1, CTIF, or MIF4GD) were loaded at 200 µM into the injection syringe. Titrations were performed using 19 injections of 2 µl each, with an injection interval of 180 s and a stirring speed of 750 rpm. Raw thermograms were processed and analyzed using a custom analysis pipeline implemented in Jupyter notebooks. Binding parameters, including binding enthalpy (ΔH), entropy (ΔS), dissociation constant (KD), and binding stoichiometry (n), were determined by fitting the data to a one-site binding model. Multiple datasets were globally fit to a single set of parameters, and 95% confidence intervals were calculated using the Fisher F-test as implemented in the lmfit Python module.

### ATPase assays

ATPase activity of human DDX19A and DDX19B and their modulation by PAIP1 were measured using an indirect NADH-coupled assay as described previously ^4^. Reactions contained, as indicated, DDX19A or DDX19B (2.5 µM), poly(A) RNA (0.1 mg ml⁻¹), and PAIP1 (10 µM). Two 2× master mixes (A and B) were prepared to enable synchronized reaction initiation. Master mix A contained proteins, BSA, glycerol, and poly(A) RNA where indicated, while master mix B contained ATP, DTT, phosphoenolpyruvate (PEP), NADH, and pyruvate kinase/lactate dehydrogenase (PK/LDH) in ATPase buffer. Equal volumes of prewarmed master mixes were combined in 384-well plates (Corning), yielding final assay conditions of 30 mM HEPES-NaOH (pH 7.5), 100 mM NaCl, 2 mM MgCl₂, 10% (w/v) glycerol, 0.33 mg ml⁻¹ BSA, 2.5 mM ATP, 10 mM DTT, 6 mM PEP, 1.2 mM NADH, and 11–25 U mL⁻¹ PK/LDH (Sigma Aldrich). ATP consumption was monitored indirectly by measuring NADH at 340 nm every minute for approximately 2 h at 37 °C using a Spark multimode plate reader (Tecan). Pathlength-corrected absorbance values were converted to NADH concentrations using the Beer–Lambert law (ε₃₄₀ = 6220 M⁻¹ cm⁻¹). Reaction rates were determined by linear regression and normalized to DDX19 concentration.

### Phylogenetic analysis and multiple sequence alignment SBM motif

Orthologous protein sequences of PAIP1 were retrieved from OrthoDB v12.1. Multiple sequence alignments were generated using MAFFT v7, and poorly aligned positions were removed using BMGE with default parameters to eliminate ambiguously aligned regions. Phylogenetic inference was performed locally using IQ-TREE v2.3.6, with automatic selection of the best-fitting amino acid substitution model. Resulting phylogenetic trees were visualized and annotated using FigTree v1.4.4, trees were midpoint-rooted for visualization. For analysis of the SBM-containing motif, the first 18 N-terminal amino acids of respective DDX19 orthologues protein sequences were retrieved from OrthoDB v12.1 and then aligned using ClustalOmega. The resulting alignment was visualized using Jalview v2.11.5.1. Orthologs were identified using PantherDB and UniProt to generate the table summarizing protein conservation across species.

### SEC-MALS

Size-exclusion chromatography coupled to multi-angle light scattering (SEC–MALS) was performed using an Agilent 1260 HPLC system equipped with a high-performance autosampler and a multi-wavelength UV–Vis absorbance detector, connected in-line to a Wyatt Technology HELEOS II 8+ multi-angle light scattering detector and an Optilab rEX differential refractive index detector. UV absorbance was monitored at 280 and 260 nm. A Superdex 200 Increase 10/300 GL column (Cytiva) was equilibrated in SEC–MALS buffer (PBS, 137 mM NaCl, 2.68 mM KCl,10.14 mM Na₂HPO₄, 1.76 mM KH₂PO₄, pH 7.5, 0.5 mM TCEP) prior to data collection and operated at a flow rate of 0.5 ml min⁻¹ to ensure stable baseline signals from all detectors. PAIP1 samples were prepared in SEC–MALS buffer and clarified by filtration through a 0.22 µm filter prior to analysis. For each sample run, 100 µL of protein at the concentration indicated in the corresponding figure was injected. Inter-detector delay volumes, band-broadening corrections, and light-scattering detector normalization were calibrated according to standard procedures using bovine serum albumin. Molecular masses, elution concentrations, and mass distributions were calculated using ASTRA software (Wyatt Technology) with a dn/dc value of 0.185 mL g⁻¹ for protein. Weight-average molar masses were determined across the elution peak following baseline subtraction.

### RNA anisotropy binding assay

Fluorescence anisotropy assays were performed using a fluorescently labeled synthetic A_18_ RNA probe labeled at the 3′ end with 6-FAM from Sigma. Protein solutions were prepared as a 4 fold master mix using SEC buffer. A twofold (2×) master mix containing 80 mM Tris–HCl (pH 7.5), 300 mM NaCl, 4 mM MgCl₂, 1 mM DTT, 20% (v/v) glycerol, and 50 nM fluorescent probe was prepared and mixed 1:1 with protein solutions for experiments containing 2 proteins or 2:1:1 with protein solution and SEC buffer for experiments containing 1 protein. Protein– probe mixtures were incubated for 1 h at 25 °C to allow equilibration prior to measurement. Measurements were performed in black, flat-bottom 384-well plates, and fluorescence anisotropy was recorded at 25 °C using a Spark multimode plate reader (Tecan). For competition experiments, defined concentrations of PABPC1 (48 nM) or PABPN1 (4 µM) were diluted 1:1 with titrated concentrations of PAIP1, yielding final concentrations of 12 nM PABPC1 or 1 µM PABPN1. Data are presented as mean values with error bars indicating standard deviation. Binding curves were analyzed using Prism 10 (GraphPad). Direct binding data were fitted using a simple binding model assuming a constant signal from the unbound species and n binding sites on a constant species (“Simple binding fit: signal from constant species, n on constant”). Competition experiments were fitted using a nonlinear regression model of inhibitor concentration versus response with a variable slope (four-parameter logistic equation).

### Human cell culture and cell lines

All cell lines were cultured in DMEM supplemented with 10% fetal calf serum and 0.1 mg mL⁻¹ penicillin–streptomycin at 37 °C in a humidified atmosphere containing 5% CO₂. Cell lines were generated using the Flp-In system (Invitrogen), resulting in stable genomic integration of pcDNA5-based tetracycline inducible expression vectors in parental HeLa cell lines containing one FRT site^5^. Monoclonal cell lines were generated following selection with 0.3 mg mL⁻¹ hygromycin B. Construct expression was adjusted to approximately endogenous levels by western blot, using 0.5 ng/mL doxycycline for PABPC1 and 1 µg/mL for DDX19A/B. No doxycycline was added for PAIP1 because leaky expression was sufficient. All cell lines are regularly tested for mycoplasma contamination.

### RNA interference

For RNAi-mediated protein depletion, cells were transfected with siRNAs using Interferin. The following siRNAs and conditions were used: siPAIP1: GCUGCAAAAGGGGAUGAAGTT, 18 nM, 48 h; siDDX19A/B: CAAGGUGUUUGUUCUGGAUTT, 9 nM, 48 h. As a negative control a pool of non-targeting siRNAs (siTOOLs, 18 nM, 48 h) was used.

### Immunostaining of fixed human cells

Cells were fixed with 3% paraformaldehyde (PFA) at room temperature for 15 min, followed by permeabilization with 0.1% Triton X-100 and 0.02% SDS in PBS for 7 min. After permeabilization, cells were washed with PBS and blocked with 2% BSA in PBS for at least 30 min. Cells were then incubated with primary antibodies diluted in 2% BSA in PBS for 1 h, followed by PBS washes and incubation with appropriate secondary antibodies. After additional washing steps with PBS, coverslips were mounted onto glass slides using Vectashield mounting medium for subsequent confocal microscopy.

### Confocal microscopy of PAIP1-mEGFP constructs

Confocal images of PAIP1-GFP and mutants were acquired on a Zeiss LSM700 upright microscope using a 63×/1.4 NA oil-immersion Plan-Apochromat objective. For all experiments, optical sections corresponding to the nuclear midplane were imaged. In panels, we show a single z plane. Images were segmented, and image features were extracted using CellProfiler.

### Polyadenylated RNA fluorescence in situ hybridization

Cells were fixed with 3% paraformaldehyde (PFA) for 10 min and washed four times with PBS. Cells were then permeabilized by incubation in 70% ethanol overnight at 4°C, followed by a wash with PBS supplemented with 10% formamide for 5 min at room temperature. Hybridization was performed overnight at 37 °C in hybridization buffer (0.5% BSA, 10% dextran sulfate, 2× SSC, 10% formamide, 200 nM yeast tRNA, and 1 ng µl⁻¹ Cy5-labeled poly(dT30) probes (Microsynth) (sequence listed in Supplemental Table 2), prepared under RNase-free conditions). Following hybridization, samples were washed three times for 30 min at room temperature with 2× SSC supplemented with 10% formamide. Coverslips were mounted onto glass slides using Vectashield mounting medium and sealed with nail polish. Nuclear and cytoplasmic intensities were classified and analyzed using CellProfiler v4.2.8.

### SplitVenus bimolecular fluorescence complementation (BiFC) assay

HeLa cells were cultured on coverslips and transiently transfected with SplitVenus constructs using lipofectamine 3000 (Invitrogen) according to the manufacturer’s protocol, using 800 ng DNA for each plasmid. Cells were incubated for 48 h post-transfection, after which expression was induced using 0.5 µg/ml doxycycline for 8h to maintain low expression and minimize false-positive rates. Cells were subsequently fixed with 3% PFA for 15 min, washed three times with PBS, and permeabilized with 0.1% Triton X-100 in PBS for 10 min. Immunostaining was performed as described above. Confocal images were collected on an Olympus IX83 microscope fitted with a Yokogawa CSU-W1 confocal scan head (50 µm disk) and a UPLX APO 60×/1.42 oil-immersion objective.

### Western blot analysis

Samples were mixed with SDS sample buffer and loaded on SDS-PAGE gradient gels. After protein separation, proteins were transferred to an activated PVDF membrane by semi-dry blotting. Membranes were blocked with 5% milk in PBST for at least 30 min, and incubated in 5% milk/PBST containing primary antibody overnight at 4°C or 2h at RT. After three washes with PBST for 5 min, membranes were incubated with a corresponding secondary antibody in 5% milk/PBST. Following three further washes with PBST, the signal was captured using Immoblon Western Chemiluninescent HRP substrate (Milipore) with a Fusion (Vilber) imaging system.

### Affinity purification for Mass Spectrometry or westernblotting

HeLa cells stably expressing protein of interest at near endogenous levels were cultured to approximately 80% confluency, washed thrice with PBS, and harvested into lysis buffer containing 0.1% Triton X-100, protease inhibitors, Benzonase, and DNase I in PBS (Benzonase was omitted for immunoprecipitations of PABPC1-mCherry). Cells were lysed by douncing, and lysates were clarified by centrifugation at 20 000 xg for 30 minutes. Clarified lysates were incubated with lysis buffer pre-equilibrated affinity beads. GFP-Trap magnetic agarose or RFP-Trap magnetic agarose beads (ChromoTek) were used for GFP- or mCherry-tagged proteins, respectively, whereas V5-conjugated beads (Biorbyt) were used for V5-tagged proteins. Lysates were incubated with GFP- or RFP-Trap beads for 2 h at 4 °C or with V5-conjugated beads overnight at 4 °C. Following incubation, Beads were washed four times with PBS containing 0.1% Triton X-100. Bound proteins were eluted in SDS sample buffer containing 60 mM Tris-HCl, pH 6.8, 10% glycerol, 2% SDS, and 180 mM β-mercaptoethanol by heating at 95 °C for 5 min. Eluates were analyzed directly by western blotting or processed for mass spectrometry as described below.

### Enhanced RNA interactome capture

HeLa cells were cultured in three conditions: control (no treatment), PAIP1 depletion by siRNA, or with PAIP1 overexpression at approximately endogenous levels. Cells were washed with ice-cold PBS and UV-crosslinked at 254 nm with an energy of 150 mJ/cm² using a UV Stratalinker 2400 (Stratagene). Cells were collected and lysed by douncing in a buffer containing 20 mM Tris-HCl, pH 7.5, 500 mM LiCl, 1 mM EDTA, 5 mM DTT, 0.5% (w/v) LiDS, and protease inhibitors. Lysates were snap-frozen in liquid nitrogen. Once all samples had been prepared, lysates were thawed in a 37 °C water bath and clarified by centrifugation at 20,000 × g for 5 min. The clarified supernatants were supplemented with DTT and incubated with pre-equilibrated oligo(dT)25 magnetic beads (New England Biolabs, cat. no. S1419S) for 30 min. Beads were washed once with lysis buffer, twice with buffer 1 containing 20 mM Tris-HCl, pH 7.5, 500 mM LiCl, 1 mM EDTA, 5 mM DTT, and 0.1% (w/v) LiDS, twice with buffer 2 containing 20 mM Tris-HCl, pH 7.5, 500 mM LiCl, 1 mM EDTA, 5 mM DTT, and 0.02% (v/v) NP-40, and twice with buffer 3 containing 20 mM Tris-HCl, pH 7.5, 200 mM LiCl, 1 mM EDTA, 5 mM DTT, and 0.02% (v/v) NP-40. Bound material was eluted by digestion with RNase T1 and RNase A. Eluates were concentrated using a SpeedVac and processed for mass spectrometry as described below.

### Preparation of samples for mass spectrometry

#### Affinity-purification samples

Affinity-purification eluates were concentrated using a SpeedVac and processed using the S-Trap sample preparation system (Protifi) according to the manufacturer’s instructions to generate peptides for LC–MS/MS analysis.

#### Enhanced RNA interactome capture samples

Concentrated RNA interactome capture eluates were processed using SP8 cleanup according to the manufacturer’s instructions. The resulting peptides were resuspended in 0.1% aqueous formic acid. For each sample, 0.3 µg of peptides was loaded onto an EvoTip by centrifugation according to the manufacturer’s instructions (Evosep, Odense, Denmark).

#### *Drosophila* embryo samples

Stage 4 and stage 7 *Drosophila* embryos from control and maternal PAIP1-depleted conditions were collected, frozen, and processed for proteomic analysis. Embryo pellets were lysed on ice in 50 µL of buffer containing 2 M guanidinium hydrochloride, 0.1 M ammonium bicarbonate, and 5 mM TCEP. Samples were homogenized using a handheld homogenizer and heated at 95 °C for 10 min. Lysates were subsequently sonicated for 10 min (Pixul). Protein concentrations were determined by tryptophan-based MPlex measurements. For each sample, 50 µg of protein was diluted with 0.1 M ammonium bicarbonate to a final guanidinium hydrochloride concentration of 0.25 M. Proteins were alkylated with chloroacetamide and digested overnight with 1 µg of trypsin. Peptides were acidified with TFA, purified using PreOmics iST solid-phase extraction cartridges, eluted, dried by vacuum centrifugation, and subsequently analyzed by mass spectrometry.

### Mass Spectrometry

#### Affinity-purification samples

Approximately 100 ng of peptides per sample were analyzed using an Orbitrap Exploris 480 mass spectrometer coupled to a Vanquish Neo HPLC system (Thermo Fisher Scientific). The analytical column was maintained at 60 °C. Peptides were separated at a flow rate of 0.2 µL/min using an in-house-packed reversed-phase HPLC column measuring 75 µm × 30 cm and containing ReproSil Saphir 100 C18 resin with a particle size of 1.5 µm (Dr. Maisch GmbH). Mobile phase A consisted of 0.1% formic acid in water, and mobile phase B consisted of 80% acetonitrile and 0.1% formic acid in water. Peptides were separated using a linear gradient from 2% to 8% mobile phase B over 5 min, from 8% to 25% over 45 min, and from 25% to 35% over 10 min. The mass spectrometer was operated in data-independent acquisition mode. MS1 spectra were acquired over an m/z range of 390– 910 at a resolution of 120,000 FWHM at m/z 200, with centroid data acquisition, automatic injection time, and a normalized AGC target of 300%. MS2 spectra were acquired using 10 m/z isolation windows, a normalized HCD collision energy of 28%, a normalized AGC target of 3,000%, a resolution of 15,000 FWHM at m/z 200, a precursor mass range of m/z 400–900, and a maximum injection time of 22 ms. Data were acquired in centroid mode. Each acquisition cycle consisted of one MS1 scan followed by 50 DIA isolation windows.

#### Enhanced RNA interactome capture samples

EvoTips containing 0.3 µg of peptides were analyzed using an Evosep One system (Evosep, Odense, Denmark) coupled to an Orbitrap Exploris 480 mass spectrometer (Thermo Fisher Scientific). The analytical column was maintained at 40 °C using a custom-built column heater. Peptides were separated using the standard 30-samples-per-day Evosep method. The mass spectrometer was operated in data-dependent acquisition mode. MS1 spectra were acquired in profile mode over an m/z range of 350–1,600 at a resolution of 120,000 FWHM at m/z 200. The AGC target was set to 1 × 10⁶ ions, and the maximum injection time was set to automatic. Each MS1 scan was followed by higher-energy collisional dissociation of the most abundant precursor ions using a maximum cycle time of 3 s and a dynamic exclusion time of 30 s. Singly charged ions, ions with charge states of six or higher, and ions with unassigned charge states were excluded from MS2 selection. MS2 spectra were acquired in centroid mode at a resolution of 15,000 FWHM at m/z 200. The target setting was set to standard, and the maximum injection time was set to automatic. The normalized collision energy was 27%, the isolation window was 1.4 m/z, and one microscan was acquired per spectrum.

#### Drosophila embryo samples

A total amount of 20 ng of Drosophila embryo peptides were analyzed in data-independent acquisition mode (DIA) using a timsTOF Ultra Mass Spectrometer (Bruker) fitted with a CaptiveSpray nano-electrospray ion source (Bruker) in combination with a Vanquish Neo liquid chromatography system (Thermo Fisher Scientific).

### Mass-spectrometry data processing and statistical analysis

#### Enhanced RNA interactome capture

Raw files from the enhanced RNA interactome capture experiments were processed using FragPipe with MSFragger. Spectra were searched against the human UniProt protein database downloaded on February 22, 2022, supplemented with commonly observed contaminants. Full tryptic specificity was required, with cleavage after lysine or arginine residues unless followed by proline, and up to two missed cleavages were permitted. Carbamidomethylation of cysteine was specified as a fixed modification, whereas methionine oxidation and protein N-terminal acetylation were specified as variable modifications. A target-decoy search strategy was used to control the protein-level false discovery rate at 1%. Search results were analyzed using the MSstats R package, version 4.13.0. Missing values were imputed using accelerated failure time model-based imputation. Differential abundance analysis was performed using limma, and pairwise p-values and q-values were calculated as implemented in MSstats.

#### Affinity-purification mass spectrometry

Raw files from affinity-purification experiments were processed using the Spectronaut directDIA workflow with standard parameters in Spectronaut version 19.0 (Biognosys). Spectra were searched against the human UniProt protein database downloaded on February 22, 2022, supplemented with commonly observed contaminants. Quantitative fragment-ion intensities were exported from Spectronaut as F.Area values and analyzed using the MSstats R package, version 4.13.0. Data were normalized using the default equalized-medians method, and missing values were imputed using accelerated failure time model-based imputation. Differential abundance analysis was performed using limma, and pairwise p-values and q-values were calculated as implemented in MSstats.

#### Drosophila embryo proteomics

Raw files from drosophila experiments were processed using a protein database search was performed using Spectronaut (Biognosys; version 19) and differential protein abundances between samples were determined by the Proteoflux software tool (Afanc. (2026). Afanc/proteoflux: Proteoflux v1.8.5 (Version v1.8.5) [Zenodo. https://doi.org/10.5281/zenodo.18640999)].

### SLAM-seq labeling, RNA isolation, library preparation, and sequencing

HeLa cells were transfected with 18 nM control siRNA or PAIP1-targeting siRNA using INTERFERin (Polyplus) according to the manufacturer’s instructions. After 24 h, cells were incubated with 12 µM 4-thiouridine (4sU) for an additional 24 h. The labeling medium was then replaced with fresh medium containing 1.2 mM unlabeled uridine (“chase”), corresponding to a 100-fold molar excess over 4sU. Cells were harvested at 0, 1, 4, and 8 h after initiation of the chase, with three biological replicates per condition and time point. Unlabeled siCTRL and siPAIP1 samples were processed in parallel as controls and for differential-expression analysis. At each time point, cells were lysed directly in TRIzol reagent, and total RNA was purified using the Direct-zol RNA Miniprep Kit (Zymo Research) according to the manufacturer’s instructions. RNA integrity was assessed by agarose gel electrophoresis. For each sample, 5 µg of total RNA was alkylated using the SLAMseq Kinetics Kit—Catabolic Kinetics Module (Lexogen) according to the manufacturer’s instructions. Libraries were prepared using the QuantSeq FWD V2 Library Prep Kit (Lexogen) and sequenced on an Illumina NovaSeq X instrument in single-read 100-bp mode.

### ONT sample prep

Total RNA was isolated in duplicate from HeLa cells treated with control siRNA or PAIP1-targeting siRNA. Cells were lysed in TRIzol reagent, and RNA was purified using the Direct-zol RNA Miniprep Kit (Zymo Research) according to the manufacturer’s instructions. For the Drosophila samples, total RNA was isolated in duplicate from stage 4 embryos with maternal PAIP1 depletion and from the corresponding control shRNA line lacking a driver. RNA was purified using the Direct-zol RNA MicroPrep Kit (Zymo Research) according to the manufacturer’s instructions. RNA integrity was assessed using an Agilent TapeStation. For each sample, 500 ng of total RNA was used to prepare full-length cDNA libraries with the Oxford Nanopore Technologies cDNA-PCR Sequencing Kit V14 (SQK-PCS114) according to the manufacturer’s instructions. Reverse transcription was followed by cDNA amplification using 14 PCR cycles with a 10-min extension time per cycle. Amplified cDNA concentration and fragment-size distributions were assessed using an Agilent 2100 Bioanalyzer. Libraries were barcoded using the Oxford Nanopore Technologies PCR Barcoding Kit V14 (SQK-PCB114-24) and multiplexed. The pooled library was loaded onto an R10.4.1 PromethION flow cell (FLO-PRO114M) at twice the recommended library input and sequenced on a PromethION instrument for 96 h. Raw nanopore signal data were subsequently re-basecalled using the final basecalling workflow and only reads generated by this post-run basecalling were used for downstream analysis.

### Custom Reference Genome and Annotation Preparation

For the human cell line (HeLa) experiments, the human GRCh38 primary assembly was used as the reference genome, downloaded from the GENCODE website^6^. To accurately quantify the expression of the transfected overexpression construct containing PAIP1, the GRCh38 genome was supplemented with the sequence of the pcDNA5-PAIP1-mEGFP plasmid. The construct’s sequence and annotations were parsed from a corresponding GenBank file of the plasmid using the “genbank_to_fasta_and_gtf” module from the “zavolab_pyutils” package (Zavolab_pyutils: This Repository Contains Python Modules Dedicated to Various Utilities for Genomic Data Analysis, like Library Size Normalization, Annotation Conversion Etc. n.d. Github. Accessed June 17, 2026. https://github.com/zavolanlab/zavolab_pyutils/tree/dev.). The coding sequences (CDSs) of the construct were converted into standard gene, transcript, and exon features and combined with the GENCODE basic annotation file. For Drosophila experiments, the reference genome (primary assembly, version BDGP6.54) and its annotation in .gtf format were downloaded from Ensembl release-115^7^.

### Differential expression and gene set enrichment analysis

Reads were aligned to the human reference genome (UCSC hg38) supplemented with the pcDNA5–PAIP1–mEGFP sequence using STAR version 2.7.10a. Default settings were used, except that up to 10 alignments per read were permitted (outFilterMultimapNmax = 10), only one alignment was reported for reads with equivalent mapping scores (outSAMmultNmax = 1) and reads without support from the splice-junction database were filtered using outFilterType = “BySJout”. Aligned reads were sorted and indexed using SAMtools version 1.11. Gene-level counts were generated using featureCounts from the Subread package version 2.0.6, applying an exon-union counting model based on Ensembl release 112 gene annotations. Count data were normalized by the trimmed mean of M-values method implemented in edgeR version 4.8.2. Genes were retained for downstream analysis if they had more than 10 reads in at least one experimental group and at least 15 reads across all samples. Principal component analysis was performed using the 25% most variable genes. Differential expression was assessed using the quasi-likelihood framework in edgeR. A negative-binomial generalized log-linear model was fitted to the count data, and dispersion estimates were moderated using empirical Bayes methods. Genes with a false discovery rate below 0.05 and an absolute log2 fold change of at least 1 were considered differentially expressed. Gene set enrichment analysis was performed using the camera function in edgeR, which accounts for inter-gene correlation. Gene sets were obtained from the Molecular Signatures Database, version 2025.1. Enrichment was tested using ranked gene-level statistics from the differential-expression analysis, with all other detected genes serving as the background. Gene sets with a false discovery rate below 0.05 were considered significantly enriched.

### SLAM-seq data processing and mRNA half-life estimation

Single-end SLAM-seq reads were processed using SlamDunk v0.4.3 within a Snakemake workflow. Reads were aligned to the hg38 genome with slamdunk map, trimming the first 12 bases of read 1 (chosen based on elevated T-to-C mismatch frequency observed at these positions during QC) and with alignment parameters (-a 4 -n 10 -ss). Alignments were filtered against exon annotation (Ensembl 112, UCSC-style chromosome names) using slamdunk filter (minimum identity 0.9), restricted to uniquely mapping reads (NH == 1) with samtools (v1.23), and coordinate-sorted and indexed. Sample-specific SNPs were called on the filtered BAMs using slamdunk snp (minimum coverage 1, minimum variant fraction 0.2) to distinguish genuine 4-thiouridine (4sU)-induced T-to-C conversions from genomic variants, and individual VCFs were merged across samples with bcftools merge (v1.23). T-to-C conversions were quantified per gene against both exon annotation using slamdunk count, supplying the merged variant calls to mask genomic SNPs, yielding per-sample conversion and coverage counts (ConversionsOnTs, CoverageOnTs, ReadCount, TcReadCount). Per-sample count tables were aggregated into master count matrices annotated with genotype, treatment, and labelling/chase time.

Aggregated counts were imported into R v4.6.1 and assembled into a SingleCellExperiment object. Only canonical Ensembl gene identifiers (ENSG) were retained, excluding LRG and other non-standard loci. Genes with fewer than 360 total reads across all samples (equivalent to an average of 10 reads per sample across 3 genotypes, 4 timepoints, and 3 replicates) were excluded prior to modelling. For each gene, converted (T-to-C) and unconverted read counts were modelled jointly using a quasi-likelihood negative binomial generalized linear model implemented in edgeR v4.11.4 (glmQLFit). Each sample was represented as a converted/unconverted count pair sharing a common, averaged library size to prevent artificial differences in normalization between the two pools. The design matrix modelled genotype-specific intercepts and genotype-by-time interaction terms for each genotype (WT, PAIP1 overexpression [OE], and PAIP1 knockdown [KD]). The fitted genotype-by-time interaction coefficient was negated to obtain a natural-log-scale decay rate, such that a positive value indicates decay, and a negative value indicates an apparent increase in transcript abundance over time. This sign convention was applied consistently throughout. mRNA half-lives were calculated from the decay rate (expressed on a per-hour basis, since the time covariate was in hours) as the natural logarithm of 2 divided by the decay rate. Half-lives were converted from hours to minutes if required. Half-life estimates were considered unreliable and set to missing if the underlying decay rate was zero or negative (indicating no decay or an apparent increase in abundance), or if the resulting half-life exceeded 16.8 hours (2.1 times the maximum 8-hour labelling timepoint). For each gene, a partial R² was calculated as an effect-size proxy for the decay term by converting single-degree-of-freedom test statistics into effect-size measures. A pseudo t-statistic was first reconstructed from the quasi-likelihood F-statistic, taking advantage of the standard statistical identity that an F-statistic with one numerator degree of freedom is equal to the square of the corresponding t-statistic. The sign of this pseudo t-statistic was taken from the sign of the original fitted slope, to preserve directionality.

The partial R² was then obtained from this pseudo t-statistic and its associated denominator degrees of freedom, using the empirical-Bayes-moderated degrees of freedom from the quasi-likelihood F-test (fit$df.total, combining the residual and prior degrees of freedom). The same degrees of freedom were used to compute the corresponding p-value. Differences in decay rate between genotypes (OE vs. WT and KD vs. WT) were assessed using glmQLFTest on the corresponding contrasts of the genotype-by-time interaction terms. Genes were considered significant at a false discovery rate of 0.05 or lower.Nanopore Sequencing Data Processing

Raw Oxford Nanopore sequencing data (POD5 format) were processed using “nanoflowz” (Nanoflowz: This Repository Contains a Set of Workflows and Tools for Processing and Analysis of ONT RNA Long-Read Sequencing Data. n.d. Github. Accessed June 17, 2026. https://github.com/zavolanlab/nanoflowz/tree/dev.), a custom Nextflow-based pipeline designed for the end-to-end analysis of Nanopore sequencing data optimized for the cDNA-PCR Sequencing kit SQK-PCB114. Briefly, the pipeline automates the following processing steps:

1. Basecalling and poly(A)-tail Profiling: Raw signal files were basecalled using “Dorado” version 1.4.0 (utilizing the dna_r10.4.1_e8.2_400bps_sup@v5.2.0 high-accuracy model) (Dorado at Release-v1.4. n.d. Github. Accessed June 17, 2026. https://github.com/nanoporetech/dorado/tree/release-v1.4.). Poly(A)-tail lengths were simultaneously estimated by Dorado, with the configuration parameter “tail_interrupt_length” set to 1 to account for possible short interruptions in homopolymeric tracts.
2. Read Alignment: Basecalled reads were aligned to the respective reference genomes using minimap2 version 2.31^8^.
3. Gene Assignment and Cleavage Site Identification: Mapped reads were assigned to their corresponding genes using “featureCounts” version 2.1.1^9^. Concurrently, cleavage and polyadenylation sites were identified using functions from the SCINPAS package^10^, UMI-deduplicated with “umi_tools” version 1.1.6^11^ and remaining reads were annotated with relevant cleavage site and motif information with newly developed python scripts.

HeLa samples (barcodes 01-04) and Drosophila samples (barcodes 07-10) were processed in separate “nanoflowz” executions. The following quality control statistics were generated by “nanoflowz”. Initial sequencing depth ranged from approximately 2.5 million to 7.0 million raw basecalled reads per sample. Alignment demonstrated good mapping efficiency, with 87-99% of reads mapping uniquely to their respective reference genomes. Subsequent pipeline steps effectively isolated biologically relevant molecules; the extraction of valid poly(A)-tailed reads with a clear cleavage-and-polyadenylation site retained 82-86% of the aligned reads. Unique Molecular Identifier (UMI) deduplication further retained 89-94% of reads, except for one HeLa sample (bioreplicate 1, PAIP1-mEGFP-transfected) that retained only 58% of reads after UMI deduplication. Following gene assignment, the final dataset yielded per-sample between 1.7 million and 4.2 million reads with single assigned gene IDs which were used for downstream expression and poly(A)-tail length analyses.

### Expression Quantification and Poly(A)-Tail Length Analysis

Following pipeline execution, tabular data containing mapping statistics, gene assignments, and poly(A)-tail lengths (pt tags) were extracted for every aligned read. For each gene, an average poly(A)-tail length was calculated. To assess gene abundance and differential expression, raw read count matrices (with genes in rows and samples in columns) were generated. Library size-normalized abundances (log-quotients) and corresponding variances were subsequently inferred using the Sanity (Sampling-Noise-Corrected Inference of Transcription Activity) method^12^ as implemented in the “zavolab_pyutils” package (pySanity). Differential expression was evaluated by calculating fold changes between condition-average log2-quotients. Statistical significance was determined via a Bayesian Wald test, which incorporates standard errors derived from both the empirical variance between biological replicates and the modeled posterior variances. To evaluate condition-specific changes in the poly(A)-tail lengths of individual genes, differential poly(A)-tail length analysis was performed using Ordinary Least Squares (OLS) regression models. For each gene, read-level or aggregated poly(A)-tail lengths were modeled as a function of the experimental condition (e.g., PAIP1 knock-down versus control) while accounting for potential bioreplicate-specific confounding effects. Because poly(A)-tail length distributions often deviate from normality and can exhibit condition-specific variances, we employed a non-parametric bootstrapping approach to assess the stability of the regression coefficients. By resampling the data across numerous iterations at a fixed subsampling rate independent of raw read abundance, empirical confidence intervals and robust p-values were derived. This approach normalizes statistical power across expression levels but limits statistical significance to transcripts with high ONT read coverage, whereas low-coverage transcripts yield p-values close to 1. This normalized the statistical power across the dynamic range of transcription, minimizing the dependency of the significance level on basal gene expression and allowing for the rigorous identification of transcripts exhibiting significant extension or shortening of their poly(A)-tails.

### Bioreplicate consistency of poly(A)-tail measurements

Reproducibility of ONT-derived poly(A)-tail length measurements was assessed using two siCTRL and two siPAIP1 biological replicates. Analysis was restricted to genes with at least 10 ONT reads in each of the four samples, yielding 5,685 genes. For each replicate, the mean poly(A)-tail length was calculated on a per-gene basis and plotted against the corresponding biological replicate. Replicate concordance was quantified by Pearson correlation across genes measured in both replicates.

### Randomization test of the global poly(A)-tail shift

To test whether the transcriptome-wide increase in poly(A)-tail length following PAIP1 depletion could be explained by sample assignment alone, we performed a within-gene label permutation test. For each gene, the four replicate-specific mean tail-length measurements (two siCTRL and two siPAIP1) were randomly reassigned to two “control” and two “knockdown” labels, and the transcriptome-wide median tail-length change (median of per-gene [siPAIP1 − siCTRL]) was recalculated. This procedure was repeated 2,000 times to generate the null distribution expected if condition labels carried no information (grey histogram; dashed line, null median ≈ 0 nt; dotted lines, 95% null interval indicated).

### Correlation of PAIP1 data with translation efficiency, NMD and other features

Transcript features were tested for association with PAIP1-dependent poly(A)-tail change and, in parallel, mRNA half-life change: steady-state abundance (log₂A, the average log₂ abundance / MA-plot ordinate from differential-expression analysis of the untreated input), translational efficiency (measured HeLa ribosome-profiling TE^13,14^), exonic (mature-transcript) length, 3′UTR length, 3′UTR GC content, 3′UTR structure (minimum free energy per nucleotide), exon count and uORF presence (Ensembl canonical transcript, GRCh38 release 111), baseline mRNA half-life and baseline poly(A)-tail length (this study), NMD sensitivity (mean abundance change on UPF1/SMG knockdown^15^), and the codon stabilization coefficient (CSC^16^). Analysis was restricted to transcripts with all features available. To guard against measurement-noise artefacts (low-abundance transcripts having noisier estimates), the response variable was the noise-normalized magnitude of change (the absolute noise-normalized z-score of the poly(A)-tail change (siPAIP1/siCTRL), and correspondingly for half-life change) rather than the raw fold-change. First, each feature’s marginal association with tail-change magnitude is shown as its Spearman rank correlation (ρ) computed one feature at a time. Second, all nine features were entered together in a multivariate rank regression (features and response rank-transformed), and the resulting regression coefficients (β) quantify each feature’s independent contribution after accounting for the others; error bars are 95% confidence intervals from 1,000 bootstrap resamples. Because candidate predictors are mutually correlated, multicollinearity was screened beforehand by pairwise correlation and rank-based variance-inflation factors (all VIF < 5). Analyses were performed in Python (pandas, numpy, scipy).

### Drosophila strains and genetics

Maternal knockdown of Paip1 was achieved using the GAL4/UAS system. The maternal driver line P{matα4-GAL-VP16}V37 (Bloomington #7063), carrying a homozygous insertion on the third chromosome, was used to drive expression of RNAi constructs during oogenesis. The following UAS-RNAi lines targeting Paip1 were used: P{TRiP.HM04067}attP2 (Bloomington #31756), P{GD13792}v26916 (VDRC #26916), and P{KK107345}VIE-260B (VDRC #110576). Virgin females carrying the maternal driver were crossed to males from each RNAi line. In the F1 generation, females carrying the maternal driver and shRNA construct, either UAS-shRNA/+; matα4-GAL-VP16/+ or matα4-GAL-VP16/UAS-RNAi, were selected and crossed with yw; t males. Embryos from these crosses were collected as described below. For control crosses, UAS-RNAi/+ females lacking the driver were selected and crossed with yw; t males.

### Embryo collection and processing

Embryos were collected from flies maintained in collection chambers on grape juice agar plates supplemented with fresh baker’s yeast and transferred daily to fresh plates. For each cross, embryos were collected for 60 min at 25 °C and aged either for 60 min (stage 4; pre-zygotic) or 180 min (stage 7; post-zygotic). Embryos were dechorionated in ∼4% sodium hypochlorite for 3 min at room temperature, washed sequentially with water, embryo wash buffer (0.7% NaCl, 0.03% Triton X-100), and again with water, and subsequently processed for mRNA extraction according to the directzol microprep protocol (Zymo).

### RT-qPCR

RT–PCR was performed using the OneStep RT–PCR Kit (Qiagen, 210212) according to the manufacturer’s instructions. Quantitative PCR (qPCR) was carried out on a Roche LightCycler II using FastStart Universal SYBR Green Master Mix (Roche, 04913914001) in 7 µL reaction volumes with a final primer concentration of 300 nM. Cycling conditions followed the manufacturer’s recommendations. Oligonucleotides are listed in Supplemental Table 2.

### Embryonic cuticle preparations

Embryos were collected on grape juice agar plates supplemented with fresh yeast at 25 °C for 12–24 h and allowed to develop for an additional 24 h. Hatched larvae were removed by repeated yeast transfers. Remaining embryos were dechorionated with 3% Chlorox for 5 min, washed, and devitellinized by vigorous shaking in methanol/heptane (1:1). Devitellinized embryos were collected, washed three times in methanol, and mounted in 73Hoyer’s medium. Preparations were spread on slides, covered with coverslips, and incubated at 58 °C for 24 h. Slides were flattened and analyzed by darkfield and phase-contrast microscopy.

### Scoring of abdominal segmentation defects in adult flies

Adult flies derived from maternal Paip1 RNAi knockdown crosses and corresponding controls were analyzed for abdominal segmentation defects. Flies were examined using a stereomicroscope (binocular), and the dorsal side of abdominal segments A1–A6 was visually inspected. Individuals were classified as either “no defect” or “with defect.” Segmentation defects were predominantly observed in segment A4 and included phenotypes such as reversed polarity, fusion with adjacent segments, partial loss, or complete absence of the segment. Control flies were scored in parallel. Only dorsal defects were systematically recorded, although subsequent inspection indicated that ventral defects were also present, suggesting that defect frequencies may be underestimated.

### Embryonic hatch rate analysis

Embryonic viability was assessed by measuring hatch rates following maternal Paip1 RNAi knockdown. Embryos were collected at low density on grape juice agar plates and incubated at 25 °C for approximately 48 hours. After incubation, embryos were scored as either “hatched” (empty eggshell) or “dead” (unhatched embryos displaying a brownish appearance). The proportion of hatched versus dead embryos was calculated for each genotype and compared to control crosses processed in parallel.

### Patterning genes network analysis

PAIP1-dependent changes were integrated into a gene-keyed table (13449 genes; FlyBase FBgn identifiers) comprising two molecular layers: poly(A)-tail length (stage-4 embryos) and protein abundance (stage-7 embryos; mass spectrometry). All measurements are reported as knockdown relative to control (PAIP1-KD/control). Significance thresholds were adjusted P < 0.05 for poly(A)-tail length and q < 0.01 for protein abundance. The anterior–posterior segmentation hierarchy was represented as a fixed, signed, directed network of 25 genes organized into four developmental tiers (maternal, gap, pair-rule, and segment-polarity) and connected by 47 curated regulatory interactions (18 activating, 29 repressing). These edges represent established regulatory relationships from the developmental genetics literature (the canonical maternal → gap → pair-rule → segment-polarity cascade) and serve as a fixed reference topology; they were not inferred from the present data. Node positions and edges were identical across all panels, with genes grouped by developmental tier and ordered consistently within each tier. Panels therefore differ only in the molecular measurement mapped onto each node. Node fill encodes the layer-specific log₂ fold-change, while detection and statistical significance were evaluated independently for each molecular layer, allowing a gene to be quantified in one panel but absent or non-significant in another.

### Statistical analysis

#### ATPase assay

Statistical significance was assessed using an ordinary one-way ANOVA with Tukey’s multiple-comparisons test. Significance for selected pairwise comparisons is indicated as follows: ns, P ≥ 0.1234 ; P < 0.0332(*); P < 0.0021(**); P < 0.0002(***); P < 0.0001(****). Experiments were performed in duplicate and repeated in three independent biological replicates.

#### Nuclear rim enrichment of PAIP1-mEGFP mutants

Cells were segmented and features extracted using Cell Profiler. Data were then plotted using R. PAIP1 NE/cytoplasm ratios were normalized to the mean ratio of the corresponding wild-type control, and significance was assessed using a one-sample t-test of the three biological replicate means against a normalized value of 1. P-values are stated in the respective figure legends.

#### Statistics sequencing and proteomics data

Statistical analyses specific to differential gene expression, gene-set enrichment, mRNA half-life estimation, differential poly(A)-tail length analysis, proteomics, permutation testing, and multivariable feature analysis are described in the corresponding Methods sections. Unless otherwise stated, statistical tests were two-sided. Comparisons between two independent groups of genes or transcripts were performed using the Mann–Whitney U test. Associations between continuous variables were assessed using Spearman rank correlation, except for biological-replicate comparisons of ONT-derived poly(A)-tail lengths and other analyses explicitly designated as Pearson correlations. Where multiple hypotheses were tested, P values were adjusted as described for the respective analysis, and a false discovery rate or adjusted P value below 0.05 was considered statistically significant unless another threshold is specified. Empirical confidence intervals for the multivariable rank-regression analysis were calculated from 1,000 bootstrap resamples. Uppercase N denotes the number of independent biological replicates, whereas lowercase n denotes the number of genes, transcripts, proteins, embryos, adult flies, or other individual observations included in an analysis. Exact sample sizes and statistical tests are provided in the figures, figure legends, or corresponding Methods sections. Statistical significance is denoted as ns, P ≥ 0.05; *P < 0.05; **P < 0.01; ***P < 0.001; and ****P < 0.0001.

#### Poly(A) RNA FISH

For the PAIP1-depletion experiment, replicate-level median nuclear-to-cytoplasmic ratios were compared using a paired two-tailed t-test. For the DDX19 depletion and rescue experiment, replicate-level median ratios were analyzed using a linear model with biological replicate included as a blocking factor, followed by Dunnett-adjusted comparisons against the siCTRL condition. P-values are stated in the respective figure legends.

