## Supplemental Table 1 for "PAIP1 couples mRNA export to cytoplasmic mRNP remodeling and poly(A) homeostasis"

| Gene | HeLa (copies/cell) | HeLa / PAIP1 | U2OS (copies/cell) | U2OS / PAIP1 | HEK293 (copies/cell) | HEK293 / PAIP1 | Liver (ppm) | Liver / PAIP1 | Whole organsim (ppm) | Whole organism / PAIP1 |
|---|---|---|---|---|---|---|---|---|---|---|
| PAIP1 | 22,976 | 1.00 | 957,604 | 1.00 | 317,838 | 1.00 | 14.2 | 1.00 | 35.2 | 1.00 |
| PABPC1 | 547,688 | 23.84 | 6,461,655 | 6.75 | 2,475,856 | 7.79 | 131 | 9.23 | 425.0 | 12.07 |
| PABPC4 | NA | NA | 1,964,842 | 2.05 | 723,517 | 2.28 | 79.1 | 5.57 | 258.0 | 7.33 |
| PABPN1 | 46,178 | 2.01 | 1,305,747 | 1.36 | 224,977 | 0.71 | 30.2 | 2.13 | 210.0 | 5.97 |
| GLE1 | 7,440 | 0.32 | 139,392 | 0.15 | 69,134 | 0.22 | 0.07 | 0.00 | 5.6 | 0.16 |
| MIF4GD | 1,960 | 0.09 | 162,296 | 0.17 | 26,320 | 0.08 | 2.85 | 0.20 | 9.9 | 0.28 |
| CTIF | 616 | 0.03 | 25,859 | 0.03 | 19,676 | 0.06 | 2.16 | 0.15 | 2.0 | 0.06 |
| DDX19A | 73,741 | 3.21 | 2,121,636 | 2.22 | 575,971 | 1.81 | 19.5 | 1.37 | 53.5 | 1.52 |
| DDX19B | 29,718 | 1.29 | 44,474 | 0.05 | 119,775 | 0.38 | 14.9 | 1.05 | 46.6 | 1.32 |

| | data source |
|---|---|
| HeLa | https://elifesciences.org/articles/16950#content |
| U2OS | https://pmc.ncbi.nlm.nih.gov/articles/PMC3261713/#S1 |
| HEK293 | https://www.science.org/doi/10.1126/science.abi6983 |
| Liver | https://pax-db.org/dataset/9606/4032974247/ |
| organism | https://pax-db.org/dataset/9606/2863052565/ |
