## Supplemental Figures for "PAIP1 couples mRNA export to cytoplasmic mRNP remodeling and poly(A) homeostasis"

Click here to access/download;Supplemental figure;SupplementalFigures.pdf

Supplemental Figures S1-S7

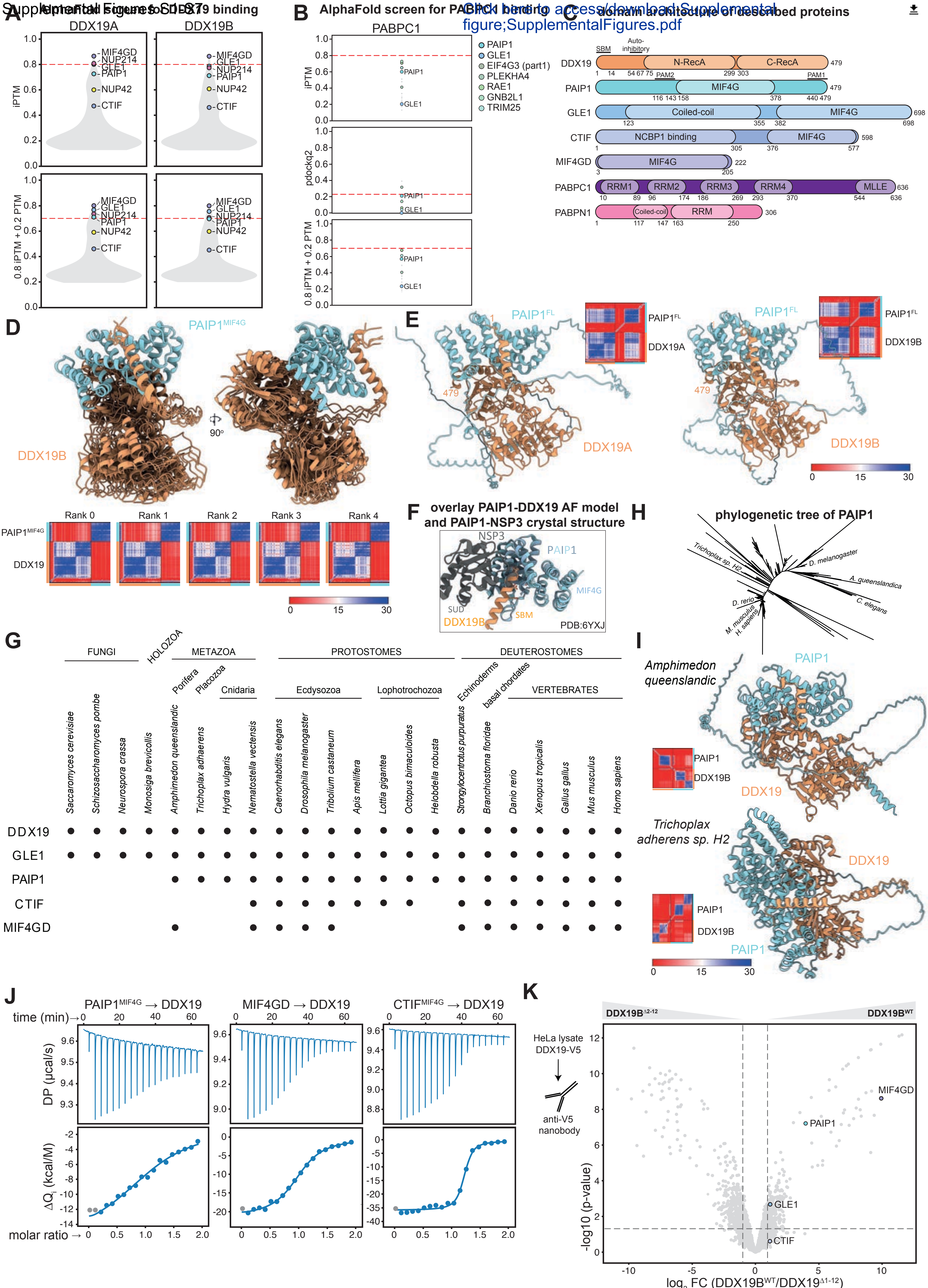


FIGURE S1

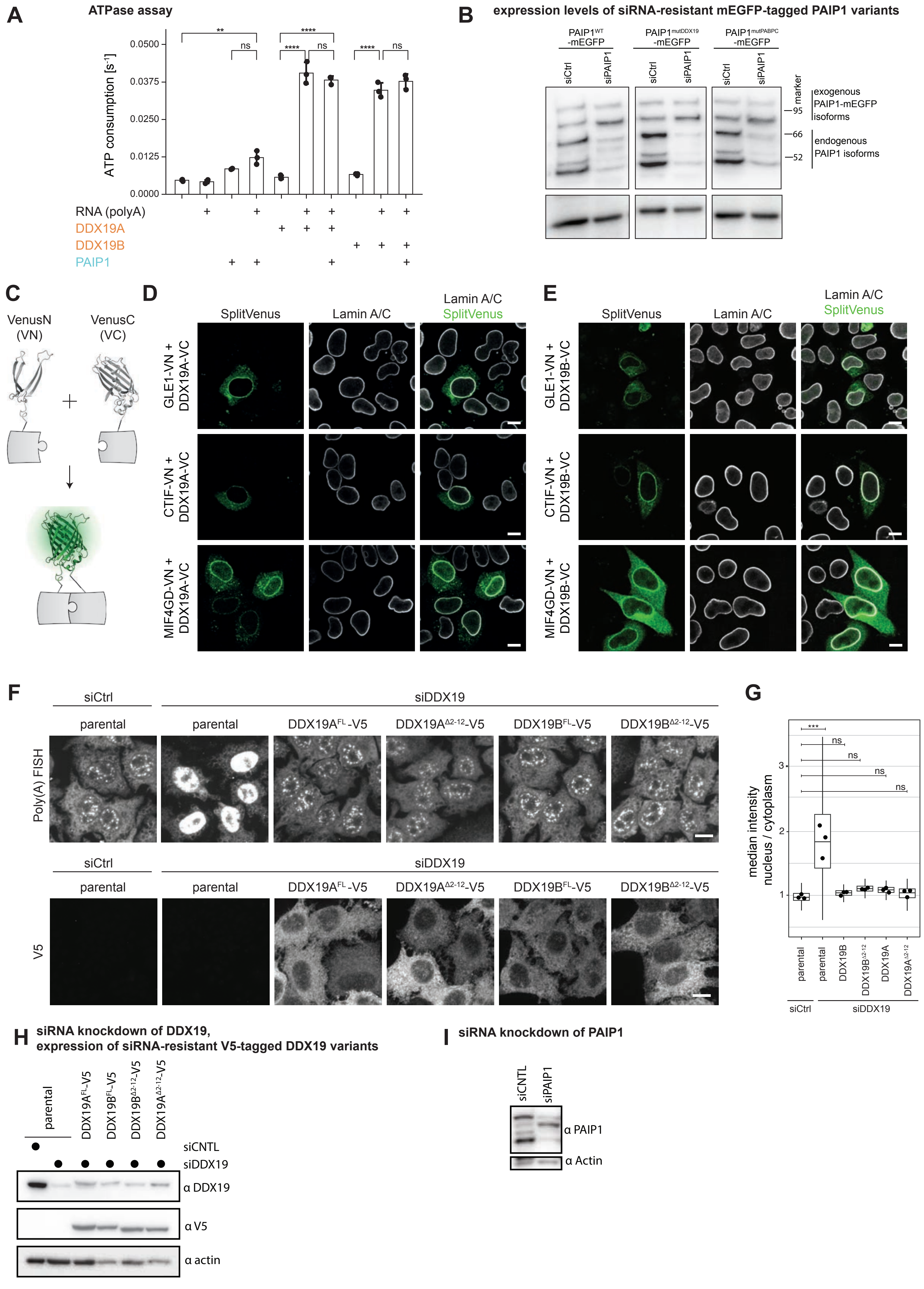
A
ATPase assay
ATP consumption [s⁻¹]
0.0500
0.0375
0.0250
0.0125
0.0000
**
****
ns
****
ns
****
ns
RNA (polyA)
DDX19A
DDX19B
PAIP1
B
expression levels of siRNA-resistant mEGFP-tagged PAIP1 variants
PAIP1WT -mEGFP
PAIP1mutDDX19 -mEGFP
PAIP1mutPABPC -mEGFP
siCtrl
siPAIP1
marker
95
66
52
exogenous PAIP1-mEGFP isoforms
endogenous PAIP1 isoforms
C
VenusN (VN)
VenusC (VC)
D
SplitVenus
Lamin A/C
GLE1-VN + DDX19A-VC
CTIF-VN + DDX19A-VC
MIF4GD-VN + DDX19A-VC
E
GLE1-VN + DDX19B-VC
CTIF-VN + DDX19B-VC
MIF4GD-VN + DDX19B-VC
F
siCtrl
siDDX19
parental
DDX19AFL-V5
DDX19AΔ2-12-V5
DDX19BFL-V5
DDX19BΔ2-12-V5
Poly(A) FISH
V5
G
median intensity nucleus / cytoplasm
***
ns
1
2
3
parental
DDX19B
DDX19BΔ2-12
DDX19A
DDX19AΔ2-12
siCtrl
siDDX19
H
siRNA knockdown of DDX19, expression of siRNA-resistant V5-tagged DDX19 variants
parental
DDX19AFL-V5
DDX19BFL-V5
DDX19BΔ2-12-V5
DDX19AΔ2-12-V5
siCNTL
siDDX19
α DDX19
α V5
α actin
I
siRNA knockdown of PAIP1
siCNTL
siPAIP1
α PAIP1
α Actin

FIGURE S2

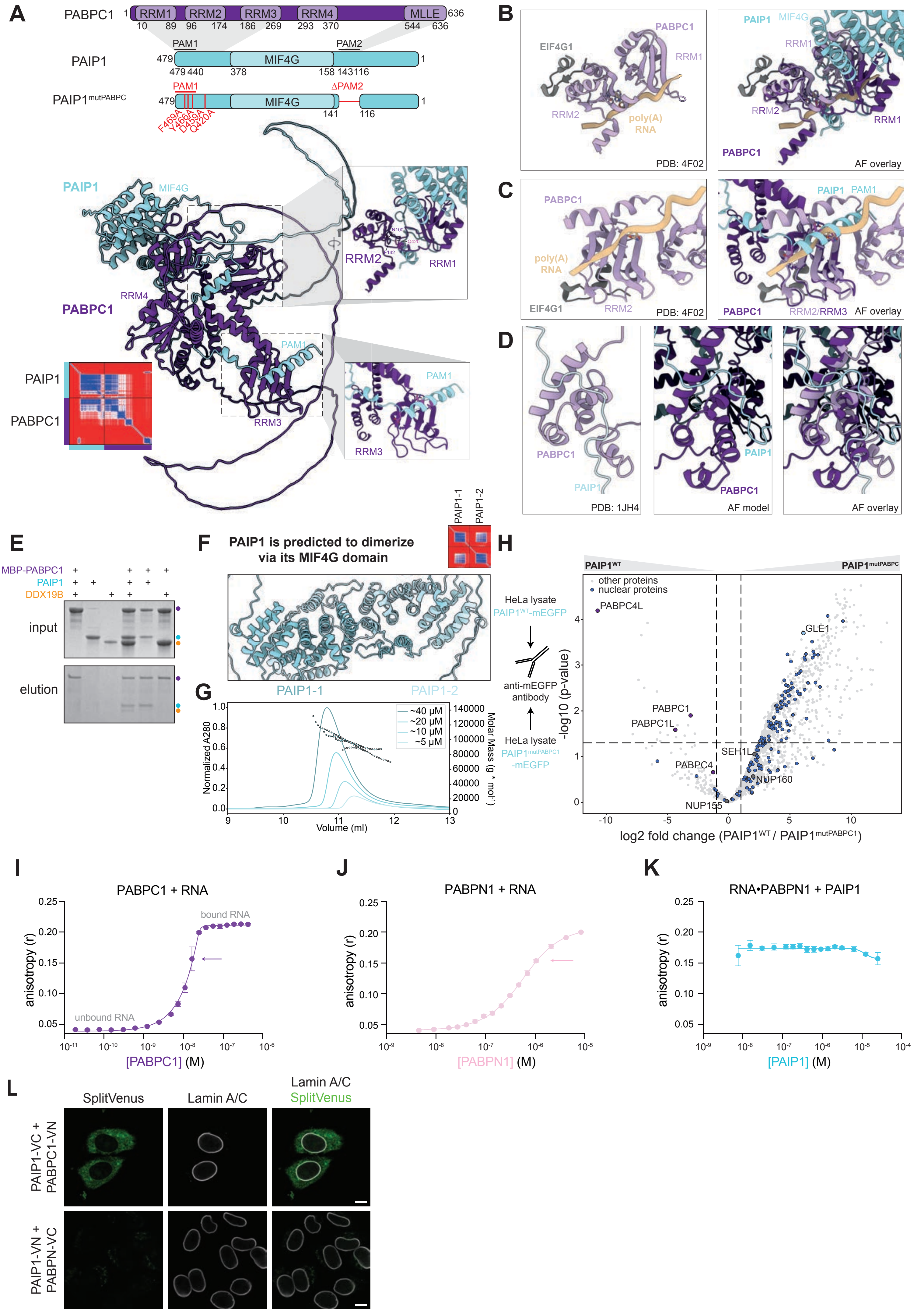
A
PABPC1
RRM1
RRM2
RRM3
RRM4
MLLE
PAIP1
PAM1
PAM2
MIF4G
PAIP1mutPABPC
ΔPAM2
F469A
Y466A
D459A
Q420A
PAIP1
MIF4G
PABPC1
RRM4
RRM3
RRM2
RRM1
PAM1
B
EIF4G1
poly(A) RNA
PDB: 4F02
AF overlay
C
D
PDB: 1JH4
AF model
E
MBP-PABPC1
PAIP1
DDX19B
input
elution
F
PAIP1 is predicted to dimerize via its MIF4G domain
PAIP1-1
PAIP1-2
G
Normalized A280
Volume (ml)
Molar Mass (g * mol⁻¹)
~40 µM
~20 µM
~10 µM
~5 µM
H
HeLa lysate
PAIP1WT-mEGFP
anti-mEGFP antibody
PAIP1mutPABPC1-mEGFP
other proteins
nuclear proteins
PABPC4L
PABPC1
PABPC1L
PABPC4
GLE1
SEH1L
NUP160
NUP155
-log10 (p-value)
log2 fold change (PAIP1WT / PAIP1mutPABPC1)
I
PABPC1 + RNA
bound RNA
unbound RNA
anisotropy (r)
[PABPC1] (M)
J
PABPN1 + RNA
[PABPN1] (M)
K
RNA•PABPN1 + PAIP1
[PAIP1] (M)
L
SplitVenus
Lamin A/C
Lamin A/C SplitVenus
PAIP1-VC + PABPC1-VN
PAIP1-VN + PABPN-VC

FIGURE S3

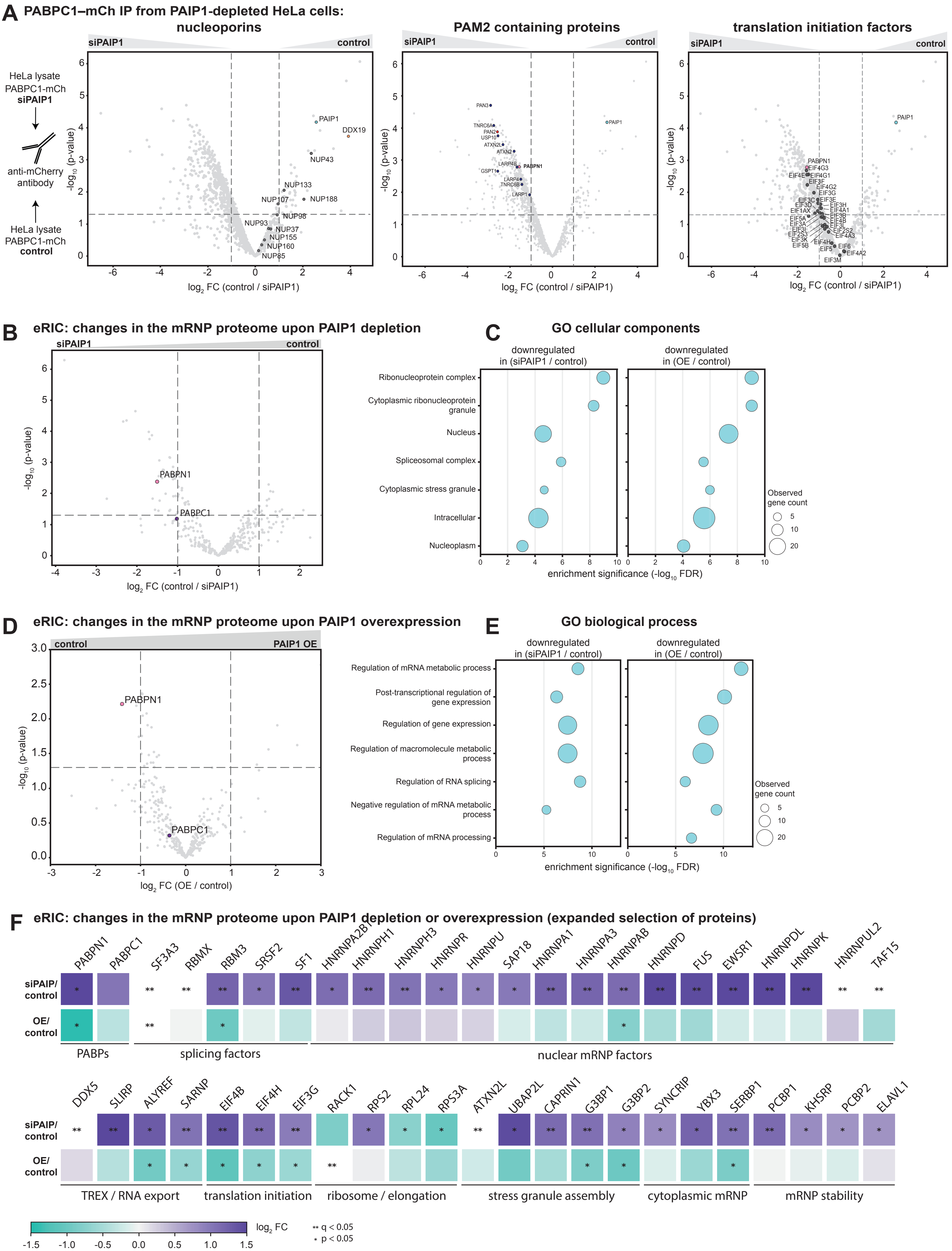
A
PABPC1–mCh IP from PAIP1-depleted HeLa cells:
nucleoporins
PAM2 containing proteins
translation initiation factors
siPAIP1
control
HeLa lysate
PABPC1-mCh
siPAIP1
anti-mCherry
antibody
HeLa lysate
PABPC1-mCh
control
-log10 (p-value)
log2 FC (control / siPAIP1)
PAIP1
DDX19
NUP43
NUP133
NUP107
NUP188
NUP98
NUP93
NUP37
NUP155
NUP160
NUP85
PAN3
TNRC6A
PAN2
USP10
ATXN2L
ATXN2
LARP4B
PABPN1
GSPT1
LARP4
TNRC6B
LARP1
EIF4G3
EIF4E
EIF4G1
EIF3F
EIF4G2
EIF3G
EIF3C
EIF3E
EIF3D
EIF3H
EIF1AX
EIF4A1
EIF5A
EIF3B
EIF2S3
EIF3A
EIF3I
EIF2S2
EIF3K
EIF4A3
EIF5B
EIF4H
EIF6
EIF5
EIF4A2
EIF3M
B
eRIC: changes in the mRNP proteome upon PAIP1 depletion
PABPC1
C
GO cellular components
downregulated
in (siPAIP1 / control)
in (OE / control)
Ribonucleoprotein complex
Cytoplasmic ribonucleoprotein granule
Nucleus
Spliceosomal complex
Cytoplasmic stress granule
Intracellular
Nucleoplasm
Observed
gene count
enrichment significance (-log10 FDR)
D
eRIC: changes in the mRNP proteome upon PAIP1 overexpression
PAIP1 OE
log2 FC (OE / control)
E
GO biological process
Regulation of mRNA metabolic process
Post-transcriptional regulation of gene expression
Regulation of gene expression
Regulation of macromolecule metabolic process
Regulation of RNA splicing
Negative regulation of mRNA metabolic process
Regulation of mRNA processing
F
eRIC: changes in the mRNP proteome upon PAIP1 depletion or overexpression (expanded selection of proteins)
PABPN1
PABPC1
SF3A3
RBMX
RBM3
SRSF2
SF1
HNRNPA2B1
HNRNPH1
HNRNPH3
HNRNPR
HNRNPU
SAP18
HNRNPA1
HNRNPA3
HNRNPAB
HNRNPD
FUS
EWSR1
HNRNPDL
HNRNPK
HNRNPUL2
TAF15
siPAIP/
control
OE/
control
PABPs
splicing factors
nuclear mRNP factors
DDX5
SLIRP
ALYREF
SARNP
EIF4B
EIF4H
EIF3G
RACK1
RPS2
RPL24
RPS3A
ATXN2L
UBAP2L
CAPRIN1
G3BP1
G3BP2
SYNCRIP
YBX3
SERBP1
PCBP1
KHSRP
PCBP2
ELAVL1
TREX / RNA export
translation initiation
ribosome / elongation
stress granule assembly
cytoplasmic mRNP
mRNP stability
log2 FC
** q < 0.05
* p < 0.05

### Figure S4

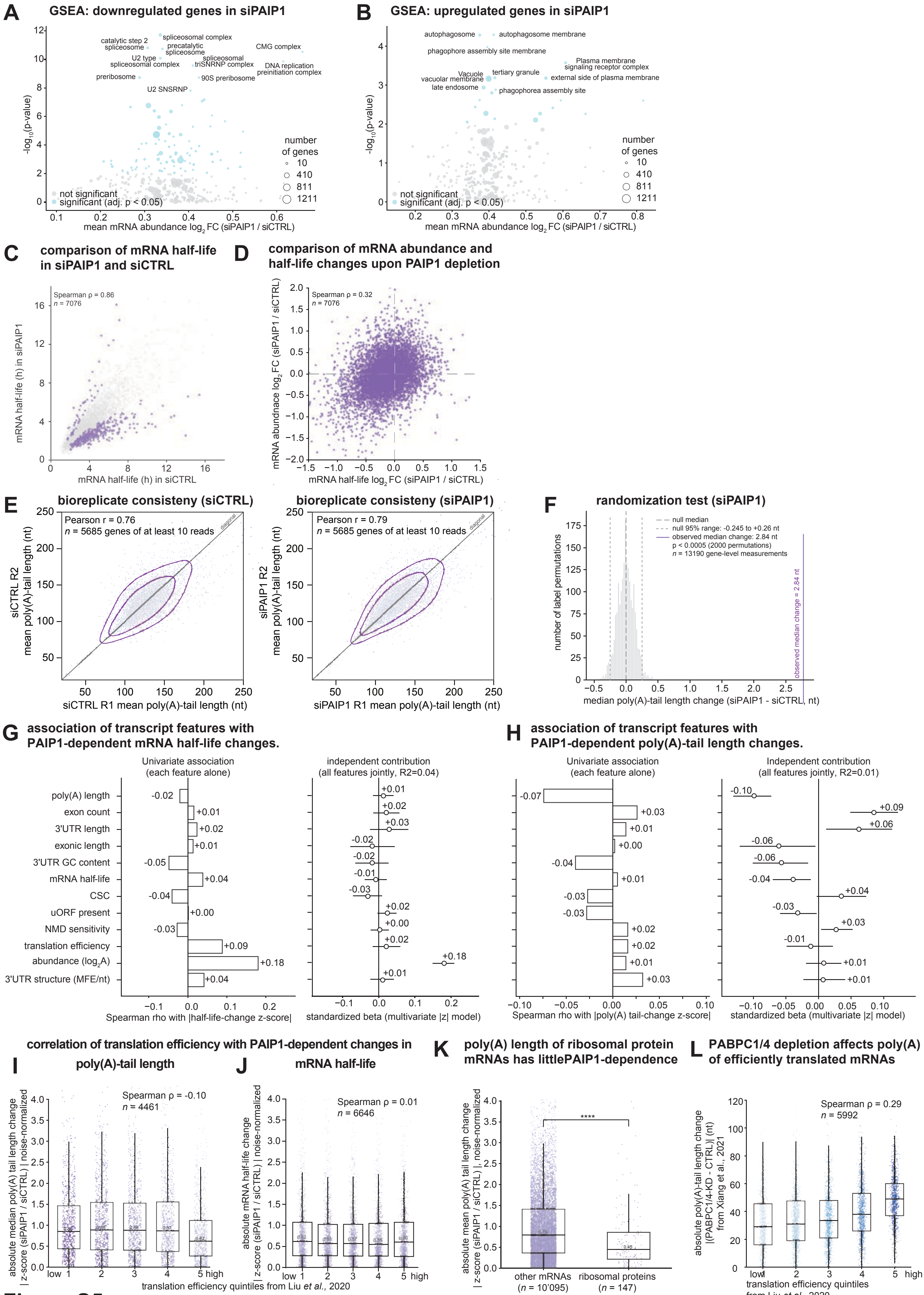
A
GSEA: downregulated genes in siPAIP1
catalytic step 2 spliceosome
spliceosomal complex
precatalytic spliceosome
CMG complex
U2 type spliceosomal complex
spliceosomal triSNRNP complex
DNA replication preinitiation complex
preribosome
90S preribosome
U2 SNSRNP
-log10(p-value)
mean mRNA abundance log2 FC (siPAIP1 / siCTRL)
number of genes
10
410
811
1211
not significant
significant (adj. p < 0.05)
B
GSEA: upregulated genes in siPAIP1
autophagosome
autophagosome membrane
phagophore assembly site membrane
Plasma membrane signaling receptor complex
Vacuole
tertiary granule
vacuolar membrane
external side of plasma membrane
late endosome
phagophorea assembly site
C
comparison of mRNA half-life in siPAIP1 and siCTRL
Spearman ρ = 0.86
n = 7076
mRNA half-life (h) in siPAIP1
mRNA half-life (h) in siCTRL
D
comparison of mRNA abundance and half-life changes upon PAIP1 depletion
Spearman ρ = 0.32
n = 7076
mRNA abundnace log2 FC (siPAIP1 / siCTRL)
mRNA half-life log2 FC (siPAIP1 / siCTRL)
E
bioreplicate consisteny (siCTRL)
Pearson r = 0.76
n = 5685 genes of at least 10 reads
diagonal
siCTRL R2 mean poly(A)-tail length (nt)
siCTRL R1 mean poly(A)-tail length (nt)
bioreplicate consisteny (siPAIP1)
Pearson r = 0.79
n = 5685 genes of at least 10 reads
siPAIP1 R2 mean poly(A)-tail length (nt)
siPAIP1 R1 mean poly(A)-tail length (nt)
F
randomization test (siPAIP1)
null median
null 95% range: -0.245 to +0.26 nt
observed median change: 2.84 nt
p < 0.0005 (2000 permutations)
n = 13190 gene-level measurements
observed median change = 2.84 nt
number of label permutations
median poly(A)-tail length change (siPAIP1 - siCTRL, nt)
G
association of transcript features with PAIP1-dependent mRNA half-life changes.
Univariate association (each feature alone)
independent contribution (all features jointly, R2=0.04)
poly(A) length
exon count
3'UTR length
exonic length
3'UTR GC content
mRNA half-life
CSC
uORF present
NMD sensitivity
translation efficiency
abundance (log2A)
3'UTR structure (MFE/nt)
Spearman rho with |half-life-change z-score|
standardized beta (multivariate |z| model)
H
association of transcript features with PAIP1-dependent poly(A)-tail length changes.
Univariate association (each feature alone)
Independent contribution (all features jointly, R2=0.01)
Spearman rho with |poly(A) tail-change z-score|
standardized beta (multivariate |z| model)
correlation of translation efficiency with PAIP1-dependent changes in
I
poly(A)-tail length
Spearman ρ = -0.10
n = 4461
absolute median poly(A) tail length change | z-score (siPAIP1 / siCTRL) | noise-normalized
J
mRNA half-life
Spearman ρ = 0.01
n = 6646
absolute mRNA half-life change | z-score (siPAIP1 / siCTRL) | noise-normalized
translation efficiency quintiles from Liu et al., 2020
K
poly(A) length of ribosomal protein mRNAs has littlePAIP1-dependence
absolute mean poly(A) tail length change | z-score (siPAIP1 / siCTRL) |, noise-normalized
other mRNAs (n = 10'095)
ribosomal proteins (n = 147)
L
PABPC1/4 depletion affects poly(A) of efficiently translated mRNAs
Spearman ρ = 0.29
n = 5992
absolute poly(A)-tail length change |(PABPC1/4-KD - CTRL)| (nt) from Xiang et al., 2021
translation efficiency quintiles from Liu et al., 2020

**Figure S5**

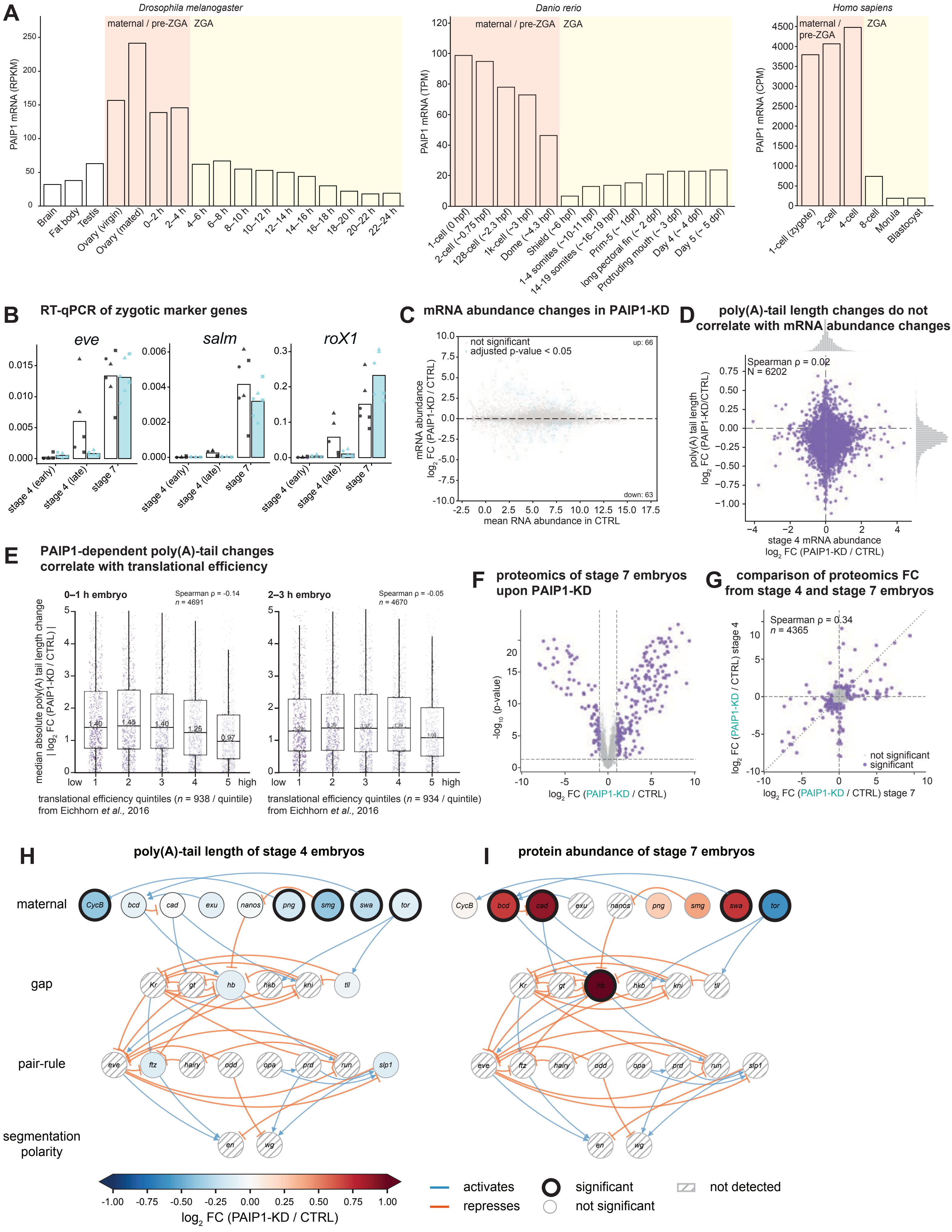
A
Drosophila melanogaster
Danio rerio
Homo sapiens
maternal / pre-ZGA
ZGA
PAIP1 mRNA (RPKM)
PAIP1 mRNA (TPM)
PAIP1 mRNA (CPM)
B
RT-qPCR of zygotic marker genes
eve
salm
roX1
stage 4 (early)
stage 4 (late)
stage 7
C
mRNA abundance changes in PAIP1-KD
not significant
adjusted p-value < 0.05
up: 66
down: 63
mean RNA abundance in CTRL
D
poly(A)-tail length changes do not correlate with mRNA abundance changes
Spearman ρ = 0.02
N = 6202
stage 4 mRNA abundance
E
PAIP1-dependent poly(A)-tail changes correlate with translational efficiency
0–1 h embryo
Spearman ρ = -0.14
n = 4691
2–3 h embryo
Spearman ρ = -0.05
n = 4670
translational efficiency quintiles (n = 938 / quintile) from Eichhorn et al., 2016
translational efficiency quintiles (n = 934 / quintile) from Eichhorn et al., 2016
F
proteomics of stage 7 embryos upon PAIP1-KD
G
comparison of proteomics FC from stage 4 and stage 7 embryos
Spearman ρ = 0.34
n = 4365
not significant
significant
H
poly(A)-tail length of stage 4 embryos
I
protein abundance of stage 7 embryos
maternal
gap
pair-rule
segmentation polarity
activates
represses
significant
not significant
not detected
log2 FC (PAIP1-KD / CTRL)

**Figure S6**

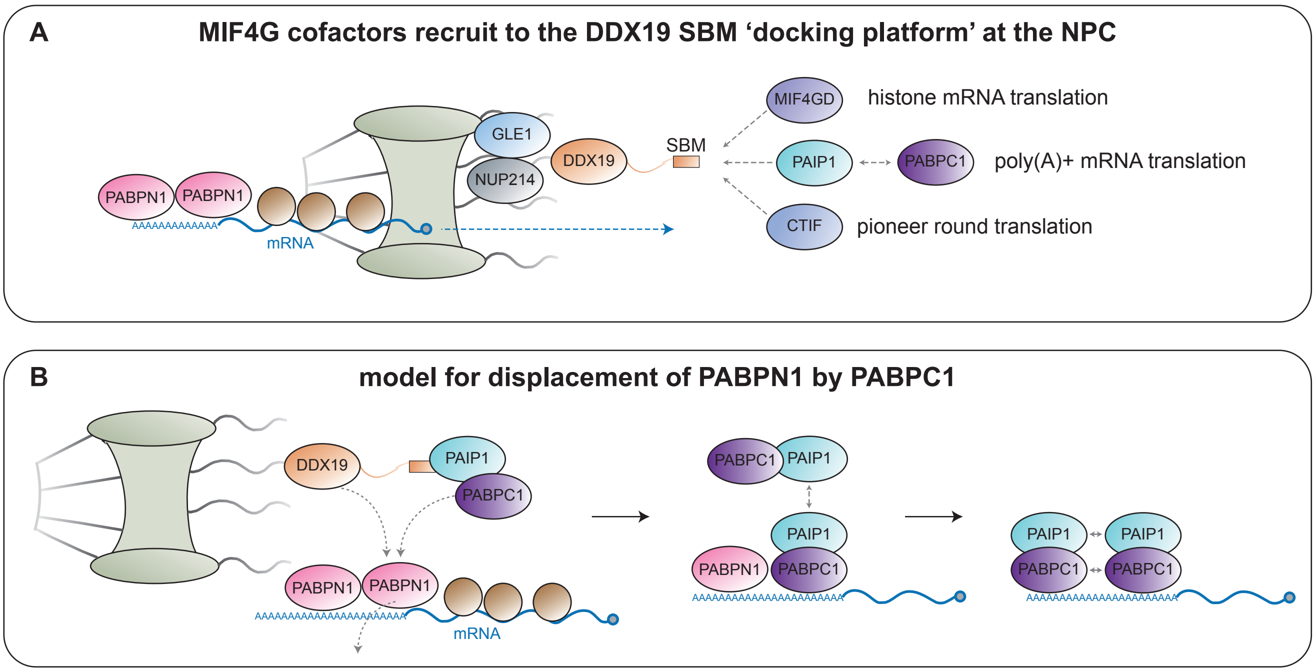
A
MIF4G cofactors recruit to the DDX19 SBM ‘docking platform’ at the NPC
MIF4GD
histone mRNA translation
GLE1
DDX19
SBM
NUP214
PAIP1
PABPC1
poly(A)+ mRNA translation
PABPN1
PABPN1
mRNA
CTIF
pioneer round translation
B
model for displacement of PABPN1 by PABPC1
DDX19
PAIP1
PABPC1
PABPN1
PABPN1
mRNA
PABPC1
PAIP1
PAIP1
PABPN1
PABPC1
PAIP1
PAIP1
PABPC1
PABPC1

FIGURE S7
